# FERDINAND: An Open Framework for Download and AI-ready Preprocessing of IMPC 2D Radiographs

**DOI:** 10.64898/2026.10.07.757287

**Authors:** Ralph Steinkamp, Holger Maier, Martin Hrabě de Angelis, Elida Schneltzer

**Affiliations:** Institute of Experimental Genetics, Helmholtz Zentrum München, Munich, Germany; German Center for Diabetes Research (DZD), Neuherberg, Germany; Chair of Experimental Genetics, School of Life Science Weihenstephan, Technische Universität München, Freising, Germany

**Keywords:** IMPC, mouse phenotyping, 2D radiographs, harmonisation, quality control, image curation, preprocessing

## Abstract

The International Mouse Phenotyping Consortium (IMPC) maintains a large archive of whole-body dorsoventral radiographs. However, heterogeneity in file formats, metadata completeness, orientation, and image quality limits its reuse for large-scale quantitative analysis. We present FERDINAND (Framework for Extracting and Restructuring Distributed IMPC radiographs into a Normalised, AI-ready Navigable Dataset), an open framework for programmatic access, harmonisation, quality control, and curation of IMPC radiographs. FERDINAND combines metadata retrieval and local radiograph archiving with centre-aware preprocessing, automated exposure screening, orientation standardisation, and clustering-assisted visual curation.

Applied to IMPC Data Release 24, FERDINAND organises 104,819 radiographs acquired between February 2012 and June 2025 across ten phenotyping centres and 6,306 single-gene knockout mouse lines, with semi-automated curation recorded for every image. As an independent validation of the harmonised output, a DeepLabCut-based bone-morphometry pipeline showed that FERDINAND-processed radiographs improve data efficiency for landmark detection: at reduced training-set sizes, they achieve better performance than original radiographs and require fewer manual annotations to reach a given level of quality.

FERDINAND provides a reproducible foundation for artificial intelligence (AI)-ready analysis and scalable quantitative phenotyping of heterogeneous IMPC and comparable radiograph resources. The complete FERDINAND source code is openly available on GitHub.

## 1 Introduction

The systematic characterisation of mouse models has become a cornerstone of functional genomics, providing mechanistic insight into gene function, development, physiology, and disease. The International Mouse Phenotyping Consortium (IMPC) [1, 2, 3] has industrialised this effort by generating and phenotyping single-gene knockout lines under standardised operating procedures, which are defined in the International Mouse Phenotyping Resource of Standardised Screens (IMPReSS) [4]. The majority of the routinely analysed IMPC output is represented as structured phenotyping parameters and processed through scalable statistical workflows [5, 6, 7]. Over the years, the IMPC programme has revealed that gene perturbation often produces phenotypes across multiple physiological systems, making the resource highly informative for the study of pleiotropy, co-morbidities, and multi-morbidities [8].

However, not all IMPC data products are already condensed into readily analysable numerical phenotypes. The consortium also generates complex and voluminous machine-derived raw data that must first be interpreted before they are transformed into statistically tractable phenotype variables. A clear example is Auditory Brainstem Response (ABR) phenotyping, where threshold identification is dependend on expert readers. Thalmeier et al. showed that supervised and self-supervised machine learning can derive objective hearing thresholds directly from ABR measurements, reducing reader dependence and opening a route towards scalable re-analysis of previously expert-only data [9]. This general problem formulation also applies to image data within the IMPC.

The IMPC pipeline includes imaging procedures, among them whole-body dorsoventral radiography [10]. In the release context targeted by this work, the corresponding image estate exceeds 100,000 radiographs and is therefore large enough to be considered a genuine data-science resource. Nevertheless, three practical gaps currently limit reuse. First, although the images are openly accessible through a well-documented application programming interface (API) and archive infrastructure [11, 12, 3], effective large-scale retrieval still requires familiarity with IMPC identifiers, procedure structure, and metadata conventions. Second, the radiographs are highly heterogeneous across centres with respect to file formats, resolution metadata, exposure, embedded labels, body position, and rotation, which makes the resource difficult to analyse as one harmonised “big data” collection. Third, current IMPC radiograph assessment is still dominated by qualitative expert review, whereas quantitative morphometry is scarcely represented in the structured database output, with `tibia_length` being the only quantitative bone metric captured [10]. Extending manual skeletal measurements to the full image resource would be prohibitively labour-intensive and costly.

Beyond replacing labour-intensive expert review, AI-based analysis can transform radiographs into a substantially richer phenotypic resource. Automated extraction of skeletal lengths, proportions, symmetry measures, and potentially more complex shape or soft-tissue features would create reproducible quantitative variables for association with genotype, sex, developmental stage, and other IMPC metadata. This would permit the existing image archive to be re-analysed at a scale that is not feasible through manual assessment and could reveal phenotypic variation that is not captured by the currently structured radiography output.

Prior work only partly addresses these challenges. In the wider IMPC ecosystem, methodological development has focused predominantly on statistical analysis of structured phenotype parameters and disease-model discovery rather than on centre-spanning image harmonisation of raw radiographs [5, 6, 7]. Automated quantitative analysis of murine radiographs has recently been demonstrated by Rong et al., who used a Keypoint R-CNN pipeline for bone-length measurements in mouse X-rays [13]. This is an important proof that quantitative skeletal readouts can be extracted automatically from radiographic images. However, the study does not provide an openly reusable end-to-end harmonisation framework for heterogeneous IMPC-style image archives, and we have not been able to identify openly available source code accompanying the publication. More broadly, landmark-based analysis of radiographs is well established in neighbouring domains such as cephalometric imaging [14], but these studies typically address anatomically narrower and technically more homogeneous image settings than multi-centre whole-body mouse radiography. A central open problem therefore remains the preparation of heterogeneous radiograph collections in a form that is robust enough for downstream large-scale AI-based analysis.

We therefore developed FERDINAND, a **F**ramework for **E**xtracting and **R**estructuring **D**istributed **I**MPC radiographs into a **N**ormalised, **A**I-ready **N**avigable **D**ataset. The framework provides reproducible download and restructuring of IMPC radiographs into a compact, consistently annotated local resource; centre-aware preprocessing and quality control that bring images into a more comparable, AI-ready form; and a foundation for downstream quantitative phenotyping. To demonstrate that this preprocessing is not only necessary but already sufficient for successful AI-based downstream analysis, we use FERDINAND-processed radiographs as input to a separate DeepLabCut-based landmark detection pipeline for automated bone morphometry [15, 16]. That morphometry workflow is not part of FERDINAND itself; rather, it serves here as an application example showing that the harmonised radiographs can support additional statistically analysable phenotypes.

This work introduces a reproducible and openly available preprocessing framework for IMPC radiographs, providing the foundation for scalable, quantitative, and community-driven image analysis in functional genomics.

## 2 Materials and Methods

### 2.1 Data

Radiographs were retrieved from the IMPC Core Data Archive (CDA) [11] through its Open Microscopy Environment (OMERO)-backed image infrastructure [17, 18]. FERDINAND was configured to assemble whole-body dorsoventral radiographs corresponding to IMPC parameter `IMPC_XRY_034_001`, scheduled for phenotyping week 14 in the Early Adult phenotyping pipeline [10].

The current dataset definition spans twelve IMPC phenotyping centres: WTSI (Wellcome Trust Sanger Institute, Hinxton, UK), JAX (The Jackson Laboratory, Bar Harbor, ME, USA), MRC Harwell (Mary Lyon Centre, MRC Harwell Institute, Harwell, UK), UC Davis (Mouse Biology Program, University of California, Davis, CA, USA), HMGU (German Mouse Clinic, Helmholtz Zentrum München, Neuherberg, Germany), CCP-IMG (Czech Centre for Phenogenomics, BIO-CEV – Institute of Molecular Genetics, Vestec, Czech Republic), TCP (Toronto Centre for Phenogenomics, Toronto, Canada), BCM (Baylor College of Medicine, Houston, TX, USA), ICS (Institut Clinique de la Souris, Illkirch, France), RBRC (RIKEN BioResource Research Center, Tsukuba, Japan), KMPC (Korea Mouse Phenotyping Center, Seoul, South Korea), and MARC (Model Animal Research Center, Nanjing University, Nanjing, China).

For each image, the metadata retrieval pipeline harmonises centre, mouse identifier, strain, date of birth, sex, cohort type, gene symbol, OMERO image identifier, acquisition date, and IMPC-reported file type into a local SQLite database [19]. Duplicate `external_sample_id/omero_id` pairs are removed during consolidation. Records labelled as `experimental` in the source metadata are standardised to `mutant`, yielding a consistent mutant-versus-control cohort definition across centres.

To match the intended phenotyping window, only radiographs acquired within *±*5 weeks of the scheduled week 14 time point are retained. In addition, records with dates of birth before 15 November 2010 are excluded in order to remove pre-IMPC legacy material from the assembled resource. Individual animals belong to one contributing centre, whereas genes may be represented across multiple centres.

FERDINAND downloads both the original archived image object and a Joint Photographic Experts Group (JPEG) derivative for each OMERO entry. In the current implementation, original files may be encountered as Digital Imaging and Communications in Medicine (DICOM), JPEG, Portable Network Graphics (PNG), or bitmap (BMP) files. Image width and height are recorded where readable for all downloaded files. Pixel-spacing metadata required for absolute millimetre-scale morphometry are recovered reliably from DICOM headers and, in a subset of PNG files, from embedded acquisition metadata, whereas JPEG files usually provide only display-oriented dots-per-inch (DPI) information. Absolute spatial calibration is therefore centre- and format-dependent and must be verified for each analysed subset. Locally, images are archived in a hierarchical directory structure organised by centre, cohort type, gene (for mutant cohorts), and sex.

### 2.2 The FERDINAND Framework

An overview of the FERDINAND workflow is shown in Figure 1. FERDINAND is implemented as a modular image-processing workflow that combines metadata retrieval, local image archiving, centre-aware preprocessing, clustering-assisted curation, automated quality control, and orientation standardisation. The operational pipeline is organised as a sequence of notebook-driven stages backed by reusable Python modules and a central SQLite database for bookkeeping. This design allows each step to be rerun independently while preserving per-image status information, derived metadata, and exclusion decisions. Although large parts of the workflow are automated, FERDINAND is intentionally not a fully hands-off pipeline. Instead, it combines scripted processing with interactive review steps, which is why the operational implementation relies on Jupyter notebooks [20] and automatically generated Hypertext Markup Language (HTML) thumbnail galleries for visual inspection and manual decision support.

**Figure 1:**
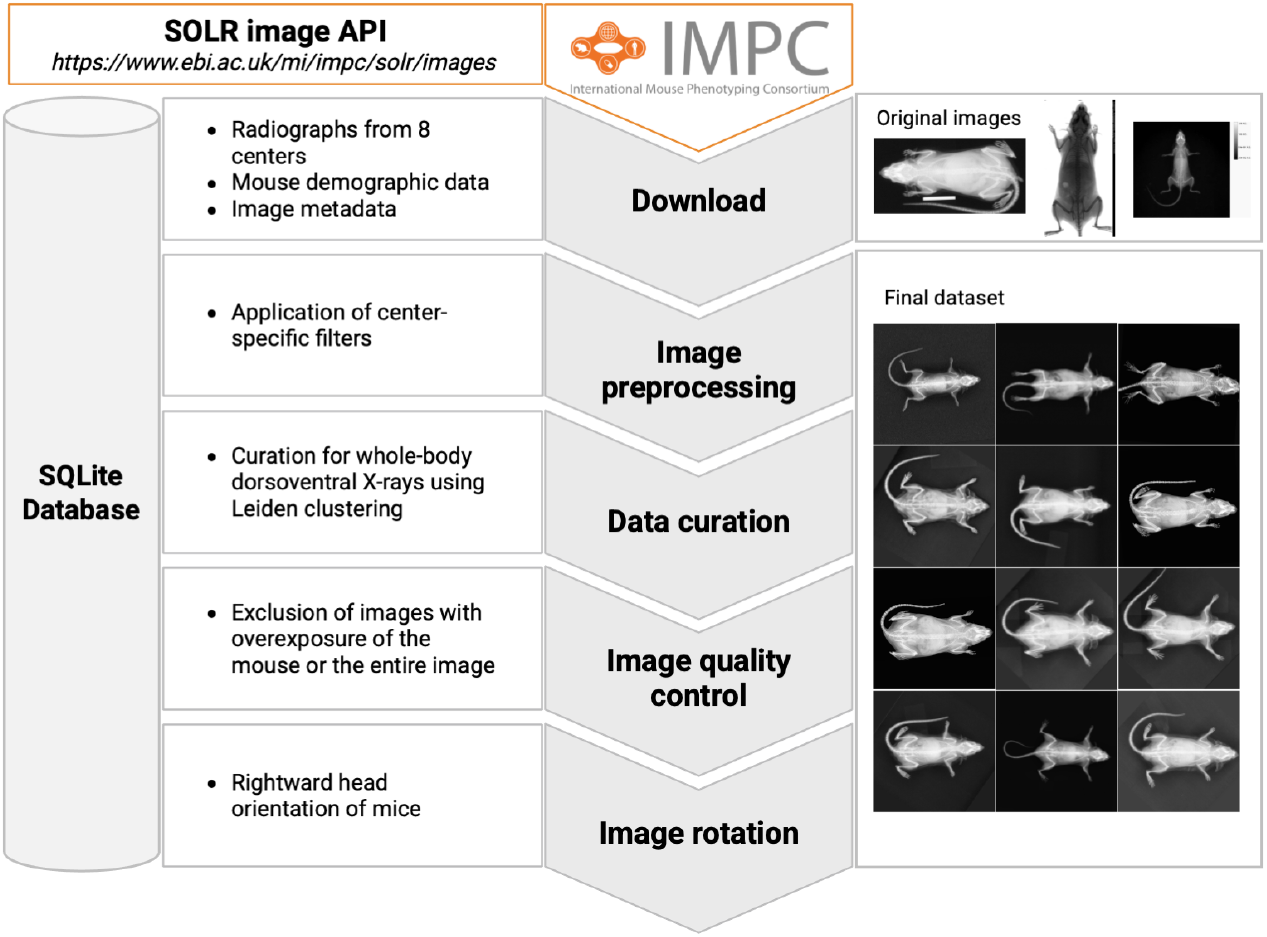
FERDINAND workflow. Radiographs and associated metadata are retrieved from the IMPC Solr image API. The SQLite database symbolised on the left records metadata and intermediate results across all processing stages. The central column shows the main workflow steps: download, centre-specific preprocessing, Leiden clustering-based curation of whole-body dorsoventral radiographs, quality control for overexposure, and rotation to a standard rightward head orientation. The right column displays representative original images and examples from the final curated dataset.

#### 2.2.1 Metadata Acquisition and Local Bookkeeping

Metadata were retrieved from the IMPC Solr image API for the configured radiography parameter (`IMPC_XRY_034_001`, whole-body dorsoventral X-ray) and for the selected phenotyping centres [12]. Listing 1 shows a complete example request.

The corresponding JavaScript Object Notation (JSON) server response is shown in Listing 2. The response illustrates the nested JSON structure and the image-level metadata ingested by FERDINAND. The production pipeline uses the equivalent request with `wt=csv` for direct tabular import; the JSON representation is shown here solely to make the response structure transparent.

**Listing 1:**
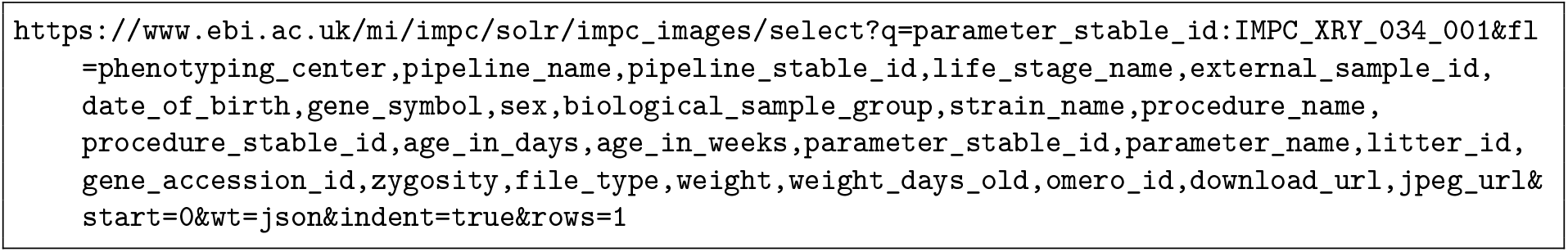
Example IMPC Solr image API request. The query filters records by the configured parameter stable identifier and requests one JSON record. Its explicit field list retrieves provenance and biological metadata, including the phenotyping centre, pipeline, specimen identifier, sex, strain, genotype, and acquisition age, together with the OMERO identifier and URLs for both the original image and its JPEG derivative.

Raw centre-specific comma-separated values (CSV) files were downloaded first and then merged into centre-level and global metadata tables. During this harmonisation step, duplicate records with the same mouse and OMERO identifier were removed, records with acquisition ages outside the configured pipeline window were excluded, and animals born before 2010-11-15 were discarded in order to restrict the dataset to the IMPC era. The metadata schema was standardised across centres, including harmonisation of mouse identifiers, cohort labels, and acquisition dates.

All consolidated metadata were stored in a SQLite database, which served as the central bookkeeping layer throughout the pipeline. The database tracked image availability, preprocessing status, image dimensions, centre-specific processing methods, curation decisions, exposure flags, orientation estimates, and rotation status. This structure made the workflow resume-safe and allowed subsequent steps to query only the images that still required processing.

#### 2.2.2 Image Download and Local Storage

For each OMERO identifier, FERDINAND downloaded the original archived image via the `download_url` metadata field and the corresponding JPEG rendering via `jpeg_url` from the IMPC image server. The original file was used to preserve format-specific metadata such as width, height, and, where available, pixel spacing; the JPEG copy provided a stable common representation for downstream image-processing steps. Image-derived metadata extracted during download were merged back into the SQLite database.

**Listing 2:**
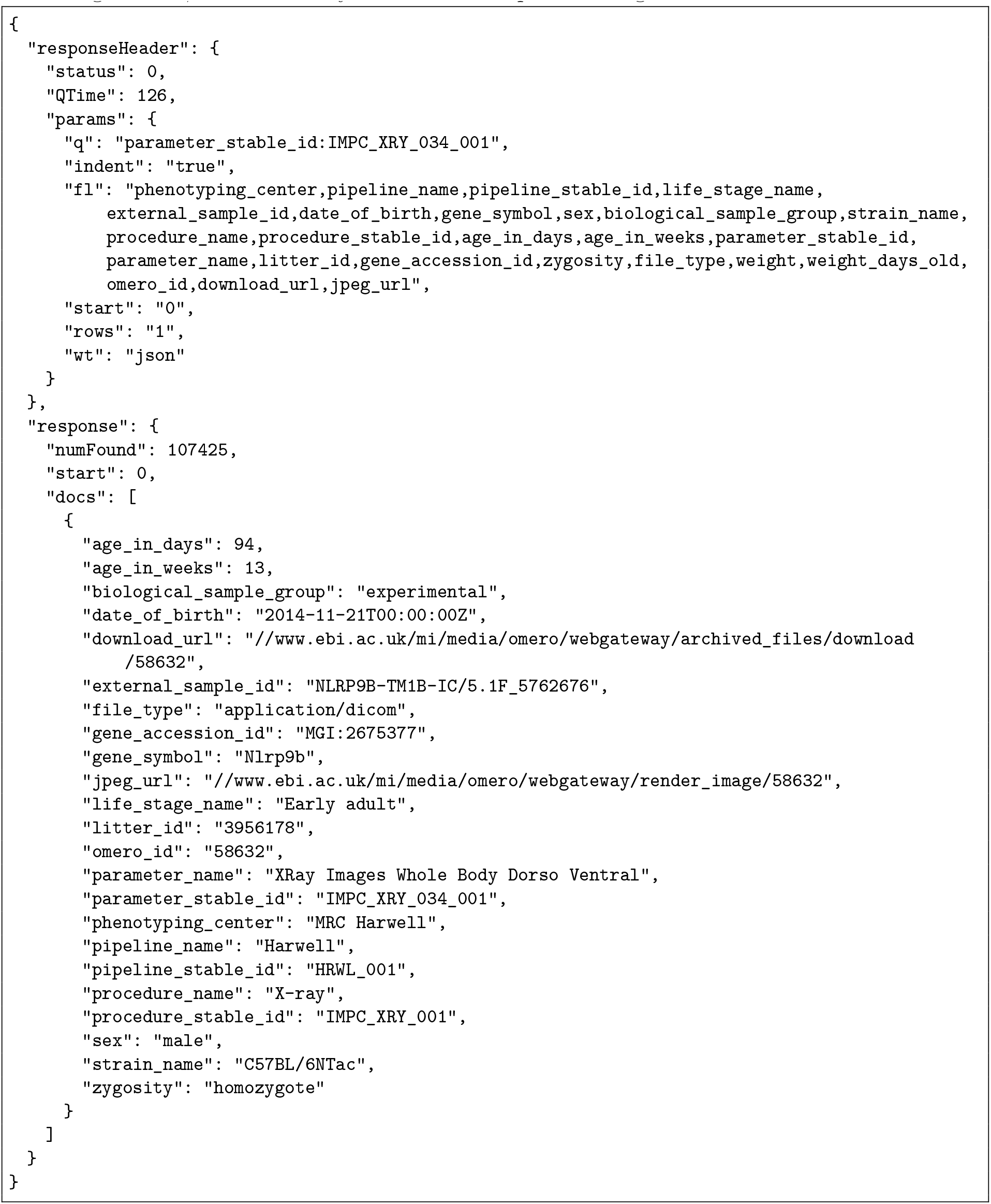
Example JSON response to the IMPC Solr image API request in Listing 1. The response header confirms the query parameters and `numFound` reports the number of matching records; the `docs` array contains the requested image metadata.

Images were stored in a hierarchical local directory structure stratified by centre, cohort type, gene symbol, and sex, with OMERO identifiers used as file names. Thumbnails and HTML galleries were generated alongside the image archive to support visual inspection at multiple stages of the workflow.

#### 2.2.3 Centre-aware Preprocessing

Preprocessing was implemented through centre-specific functions reflecting the heterogeneity of acquisition systems and image annotations across IMPC sites. A common baseline consisted of Gaussian blurring, edge enhancement, and contrast limited adaptive histogram equalisation (CLAHE) [21]. Centre-specific branches then applied operations such as cropping of text-labelled margins, inverting white-background images, removing markers, and normalising brightness.

In the current codebase, explicit preprocessing functions are implemented for BCM, HMGU, ICS, JAX, KMPC, MRC Harwell, RBRC, TCP, UC Davis, and WTSI. Examples of centre-specific rules include cropping from the right margin for BCM and bright TCP images, inversion after detecting inverted contrast for BCM and ICS, brightness-dependent marker removal for MRC Harwell, and brightness enhancement for dim UC Davis images. For centres with an implemented default branch and no additional special handling, the standard blur-edge-CLAHE sequence was applied. After preprocessing, images were centred on a square canvas using centre-dependent bounding-box- or centroid-based centring to facilitate downstream feature extraction and rotation. The exact operations applied to each image were recorded in the SQLite database as per-image processing flags. These implemented branches cover all centres represented in the current local FERDINAND database, although preprocessing support is not yet complete across the broader configured 12-centre target set.

#### 2.2.4 Clustering-assisted Dataset Curation

To identify non-standard radiographs, FERDINAND performed clustering-assisted curation on the preprocessed image set. Deep feature embeddings were extracted from red-green-blue (RGB)-converted radiographs using a pretrained vision transformer backend; the default implementation used `microsoft/swin-tiny-patch4-window7-224` [22]. Embeddings were Euclidean *L*_2_-normalised and used to build centre-specific symmetrised *k*-nearest-neighbour graphs.

Community detection was then performed with the Leiden algorithm using the Reichardt–Bornholdt configuration vertex partition [23, 24]. The resolution parameter and neighbourhood size were optimised per centre, with modularity used to guide selection of suitable clustering granularity. Cluster structure was visualised by principal component analysis (PCA) and uniform manifold approximation and projection (UMAP) [25, 26]. Interactive HTML thumbnail galleries were generated for manual review. This manual review step is essential in the current workflow: based on the galleries, entire clusters or individual images were marked for exclusion in centre-specific exclusion dictionaries and then written back to the SQLite database. The curation stage was purposefully designed as a human-in-the-loop QC step to remove lateral projections, close-ups, partial-body views, and other visually aberrant acquisitions before downstream analysis.

#### 2.2.5 Automated Exposure Quality Control (QC)

After curation, FERDINAND applied an automated exposure screen to retained images. The implemented heuristics combined a mouse-focused brightness measure with a whole-image overexposure check and wrote two explicit flags (`overexposed` and `overall_overexposed`) to the SQLite database. The notebook workflow supports an initial random-sample inspection before dataset-wide execution, reflecting that this stage is intended as a screening tool rather than a fully annotation-free replacement for visual review.

#### 2.2.6 Orientation Standardisation and Rotation Quality Control

Orientation harmonisation was performed on curated, non-overexposed images with a pretrained convolutional neural network that predicts one of 360 angular classes from a resized greyscale image. The model architecture implemented in FERDINAND consists of stacked convolutional blocks with residual connections, global average pooling, and dense classification layers [27]. For each image, a first angle prediction was stored in the SQLite database and converted to a rotation angle by taking the negative predicted orientation modulo 360.

Images were then rotated accordingly and written to a dedicated output archive. Background fill values for the rotated canvas were derived from border pixels of the source image to reduce artefactual padding contrast. A second angle prediction on the rotated images served as rotation QC. In the notebook logic, images with residual deviations from the target orientation were summarised per centre, allowing failed or imperfect alignments to be quantified and reviewed.

### 2.3 Application Example: Automated Bone Morphometry

To demonstrate downstream utility of FERDINAND, we additionally analysed radiographs with a separate DeepLabCut-based landmark detection pipeline for automated bone morphometry. In its current form, this pipeline explicitly compares landmark detection results obtained from original and FERDINAND-processed radiographs in order to quantify the benefit of FERDINAND preprocessing for downstream landmark detection. The implementation followed the standard DeepLabCut workflow described in the official documentation and primary publications [28, 15, 16] and used DeepLabCut 3.0.0 with the PyTorch engine [29]. A set of 360 distinct radiographs was annotated for 17 landmarks, defined by an expert from the German Mouse Clinic: four skull landmarks, paired proximal and distal landmarks for the left and right humerus, femur, and tibia, and one additional caudal reference landmark. The original-radiograph and FERDINAND-processed-radiograph variants were treated as two independently annotated DeepLabCut projects with separate ground-truth files rather than as a single annotation set transformed between coordinate systems. Comparisons between both image domains were therefore paired by image identity, using the same 360 source radiographs, rather than by coordinate back-transformation. The current evaluation workflow is based primarily on five-fold cross-validation rather than on a single fixed train/test split.

Because DeepLabCut expects a flat project-specific directory layout, the required folder structure was generated by scripts that created symbolic links to the corresponding radiographs. In the current pipeline, models were trained and evaluated on two image domains: original radiographs before FERDINAND processing and FERDINAND-processed radiographs. This comparison was used to test whether FERDINAND processing improves landmark-detection accuracy and data efficiency. Model training was performed with the PyTorch DeepLabCut engine and a ResNet-101 backbone. The training configuration used `TrainingFraction`=0.8, `pcutoff`=0.6, AdamW optimisation [30] with an initial learning rate of 5 × 10^−4^, a batch size of 2, and a fixed seed of 42, and RGB input conversion with image normalisation. Fold assignments were fixed in project-specific `cv_splits.csv` files and not regenerated between runs. The configured augmentation scheme combined random affine transformations (probability 0.5, rotation up to 10°, scaling 0.9-1.1), horizontal flipping with left-right landmark remapping (probability 0.5), keypoint-centred random crops of 1800 × 1800 pixels with a maximum shift of 5%, and additive Gaussian noise. Folds were generated by random shuffling of the annotated pool and were therefore not explicitly stratified by phenotyping centre.

Cross-validation performance was quantified on held-out test images by the Euclidean pixel distance between predicted and manually annotated landmarks and aggregated per image, fold, centre, anatomical group, and keypoint. In addition, DeepLabCut’s own per-fold train and test root mean squared error (RMSE), RMSE at the probability cutoff, mean average precision (mAP), and mean average recall (mAR) were recorded from the evaluation output. Percentage of correct keypoints (PCK), defined as the proportion of predictions within a specified pixel distance of the manual annotation, was evaluated at 5- and 10-pixel thresholds. Six images lacking a valid `skull_1` annotation were excluded from `skull_1`-specific error calculations only, while all remaining landmarks from these images were retained.

Beyond the main five-fold cross-validation analysis, the pipeline also included a learning-curve evaluation to quantify data-efficiency effects across reduced training-set sizes. For each of the five folds, additional models were trained at effective train fractions of 30% and 60% of the total dataset, with training images independently subsampled from the fold’s training pool. The corresponding 80% training-fraction results were reused from the main cross-validation project, since the held-out test sets were identical across all three fractions. For deployment, the five FERDINAND-processed cross-validation shuffles were further compared by a ground-truth-free quality control workflow based on inference over large per-centre image samples of up to 3000 radiographs per centre. An arbitrary ranking score combined the fraction of predictions above the DeepLabCut probability cutoff (50%), bone-length consistency (20%), failure rate (10%) and bilateral (left/right) symmetry (20%). Bone-length consistency (coefficient of variation of predicted bone length) was computed on a single, fixed set of images common to all five shuffles – namely those images for which every shuffle’s prediction cleared the probability cutoff – rather than on each shuffle’s own, separately filtered set of images. This avoids a bias whereby a shuffle with a lower overall detection rate would be scored only on an easier, self-selected subset of images and could thereby appear more consistent than a shuffle that succeeded on a larger, more difficult set of images. Cross-centre variance was recorded as an additional QC descriptor.

Paired original-versus-processed comparisons were evaluated with two-sided Wilcoxon signed-rank tests [31]. Binary percentage-of-correct-keypoints comparisonswere evaluated with exact McNemar tests [32] at the 5- and 10-pixel thresholds. Outlier sensitivity analyses of per-image paired error differences used the modified z-score criterion [33]. Left-right agreement of the resulting cleaned bone lengths was assessed by Bland-Altman analysis. Training and inference were executed in a multi-stage workflow comprising cross-validation, learning-curve analysis, deployment-model selection, and large-scale production inference. The final production inference used a FERDINAND-processed deployment model selected from the comparative evaluation workflow. The resulting landmark predictions were stored in a dedicated SQLite `bone_landmarks` table and used to derive Euclidean distances between predefined point pairs corresponding to skull length and width, as well as left and right humerus, femur, and tibia lengths. Downstream morphometric measurements were then analysed together with image identifiers and centre information from the same SQLite resource. They were further cleared by four anatomical plausibility checks (left-right symmetry via Bland-Altman analysis, femur-tibia joint-distance plausibility, skull length/width plausibility, and landmark-likelihood filtering).

The full high-throughput murine bone morphometry workflow, including model development and validation, will be described in a dedicated manuscript currently in preparation.

## 3 Results

### 3.1 The Harmonised Radiograph Resource

The current FERDINAND database comprises 104,819 whole-body radiographs from IMPC Data Release 24 [34], acquired between February 2012 and June 2025. The resource represents 6,306 single-gene knockout lines across ten phenotyping centres. The two additional configured centres, CCP-IMG and MARC, supplied no whole-body dorsoventral radiographs for the selected IMPC parameter and are therefore not represented in the local resource. At this scale, the resource already combines substantial biological diversity, centre-specific acquisition practice, and technical heterogeneity in image format and orientation, making direct downstream use without harmonisation impractical.

The composition of the assembled resource already reflects the broad experimental scope of the IMPC archive. Mutant animals account for the majority of radiographs in both sexes, while controls remain sufficiently represented to support comparative downstream analyses, as detailed in Table 1. The archived radiographs were acquired using X-ray systems from multiple manufacturer/model combinations across contributing centres (Table 3). Taken together, these summaries show that FERDINAND is not operating on a narrowly defined single-centre collection, but on a distributed multi-centre archive that already captures the diversity expected from the IMPC radiography resource.

**Table 1:** Radiograph counts by sex and cohort type, with percentages relative to the full FERDINAND resource.

| Sex | Control |  | Mutant |  | Total |  |
| --- | --- | --- | --- | --- | --- | --- |
|  | n | % | n | % | n | % |
| Female | 9,416 | 9.0 | 43,099 | 41.1 | 52,515 | 50.1 |
| Male | 9,299 | 8.9 | 43,005 | 41.0 | 52,304 | 49.9 |
| Total | 18,715 | 17.9 | 86,104 | 82.1 | 104,819 | 100.0 |

Substantial technical heterogeneity is already visible at the archive level. The image volume is distributed unevenly across centres, and the original archive spans several file types rather than a single uniform source representation, as illustrated in Figure 2. This imbalance would risk confounding pooled analyses if centre-specific imaging characteristics were ignored. Exact file-type counts, pixel-spacing coverage, and original colour-scheme metadata are reported in Table 2.

**Figure 2:**
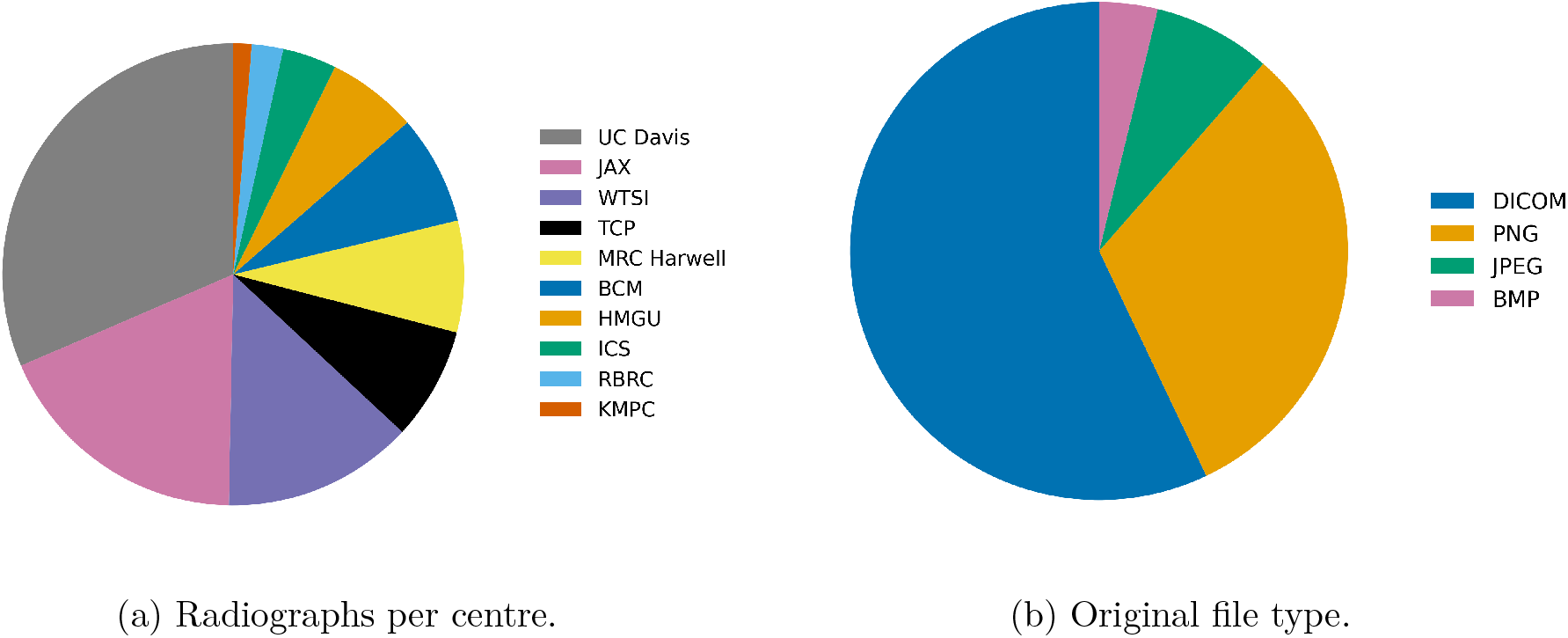
Composition of the FERDINAND radiograph resource across phenotyping centres and original archived file types.

**Table 2:** Technical characteristics of the original files in the current FERDINAND resource. Pixel-spacing eligibility is limited to DICOM and PNG; complete metadata are available for 94.7% of eligible files.

| Category | Characteristic | Count | % of resource |
| --- | --- | --- | --- |
| Original file type | DICOM | 59,836 | 57.1 |
|  | PNG | 32,981 | 31.5 |
|  | JPEG | 8,033 | 7.7 |
|  | BMP | 3,969 | 3.8 |
| Pixel-spacing metadata | Eligible formats (DICOM or PNG) | 92,817 | 88.5 |
|  | Complete row- and column-spacing meta-data (94.7% of eligible files) | 87,895 | 83.9 |
| Original colour scheme | Greyscale | 99,417 | 94.8 |
|  | Colour-coded | 5,402 | 5.2 |

**Table 3:** X-ray system manufacturer/model combinations represented in the current FERDINAND resource and the corresponding contributing centres.

| X-ray system manufacturer/model | Centres |
| --- | --- |
| Bioptics Inc / MX-20 | TCP |
| DALSA Life Sciences, Tucson, Arizona, USA / DUAL WIDE Camera | MRC Harwell, TCP |
| Faxitron Bioptics LLC / MX-20 | TCP |
| Faxitron Bioptics LLC / PathVision | HMGU, JAX, KMPC, MRC Harwell, TCP, UC Davis |
| Faxitron X-ray Corporation, Wheeling, Illinois, USA / DUAL WIDE Camera | MRC Harwell |
| Faxitron X-Ray LLC / MX-20 | HMGU, JAX, KMPC, RBRC, UC Davis, WTSI |
| Faxitron X-Ray LLC / Unknown | HMGU, JAX, KMPC, RBRC, UC Davis |
| Unknown / Unknown | BCM, HMGU, ICS, MRC Harwell, UC Davis |

Image geometry shows a similarly concentrated but non-uniform pattern. The 27 distinct original image geometries represented in the current database are condensed in Table 4 to the five most frequent layouts plus an ‘Others’ category for the remaining long tail. The full geometry distribution is provided in Supplementary Table S1.

**Table 4:** Most frequent original image geometries represented in the current FERDINAND resource. The full list is provided in Supplementary Table S1.

| Width | Height | Count |
| --- | --- | --- |
| 2368 | 2340 | 47,083 |
| 3072 | 2048 | 38,306 |
| 2048 | 1024 | 7,173 |
| 2364 | 2048 | 3,694 |
| 2048 | 2048 | 2,702 |
| Others |  | 5,861 |

The implemented preprocessing logic mirrors this site-specific heterogeneity. Gaussian blurring, edge enhancement and CLAHE form the common baseline across all centres represented in the current local database, whereas cropping, inversion, marker removal and brightness adjustment are only required for specific acquisition contexts, as summarised in Table 5. This pattern is consistent with the practical role of FERDINAND: the framework standardises a shared downstream representation while preserving sufficient flexibility to address site-specific artefacts when necessary.

**Table 5:** Centre-specific preprocessing steps implemented in the current FERDINAND dataset.

| Preprocessing step | BCM | HMGU | ICS | JAX | KMPC | MRC Harwell | RBRC | TCP | UC Davis | WTSI |
| --- | --- | --- | --- | --- | --- | --- | --- | --- | --- | --- |
| Gaussian blur | ✓ | ✓ | ✓ | ✓ | ✓ | ✓ | ✓ | ✓ | ✓ | ✓ |
| Edge enhancement | ✓ | ✓ | ✓ | ✓ | ✓ | ✓ | ✓ | ✓ | ✓ | ✓ |
| Crop right margin | ✓ |  |  |  |  |  |  | ✓ |  |  |
| CLAHE | ✓ | ✓ | ✓ | ✓ | ✓ | ✓ | ✓ | ✓ | ✓ | ✓ |
| Crop top margin |  |  | ✓ |  |  |  |  |  |  |  |
| Invert image | ✓ |  | ✓ |  |  |  |  |  |  |  |
| Remove marker |  |  |  |  |  | ✓ |  |  |  |  |
| Adjust brightness |  |  |  |  |  |  |  |  | ✓ |  |

Variation in image orientation adds a further layer of inconsistency to the raw archive. The broad pre-rotation angular distribution shown in Figure 3 confirms that orientation normalisation is not a cosmetic post-processing step, but a prerequisite for any subsequent landmark-based or other geometry-sensitive analysis.

**Figure 3:**
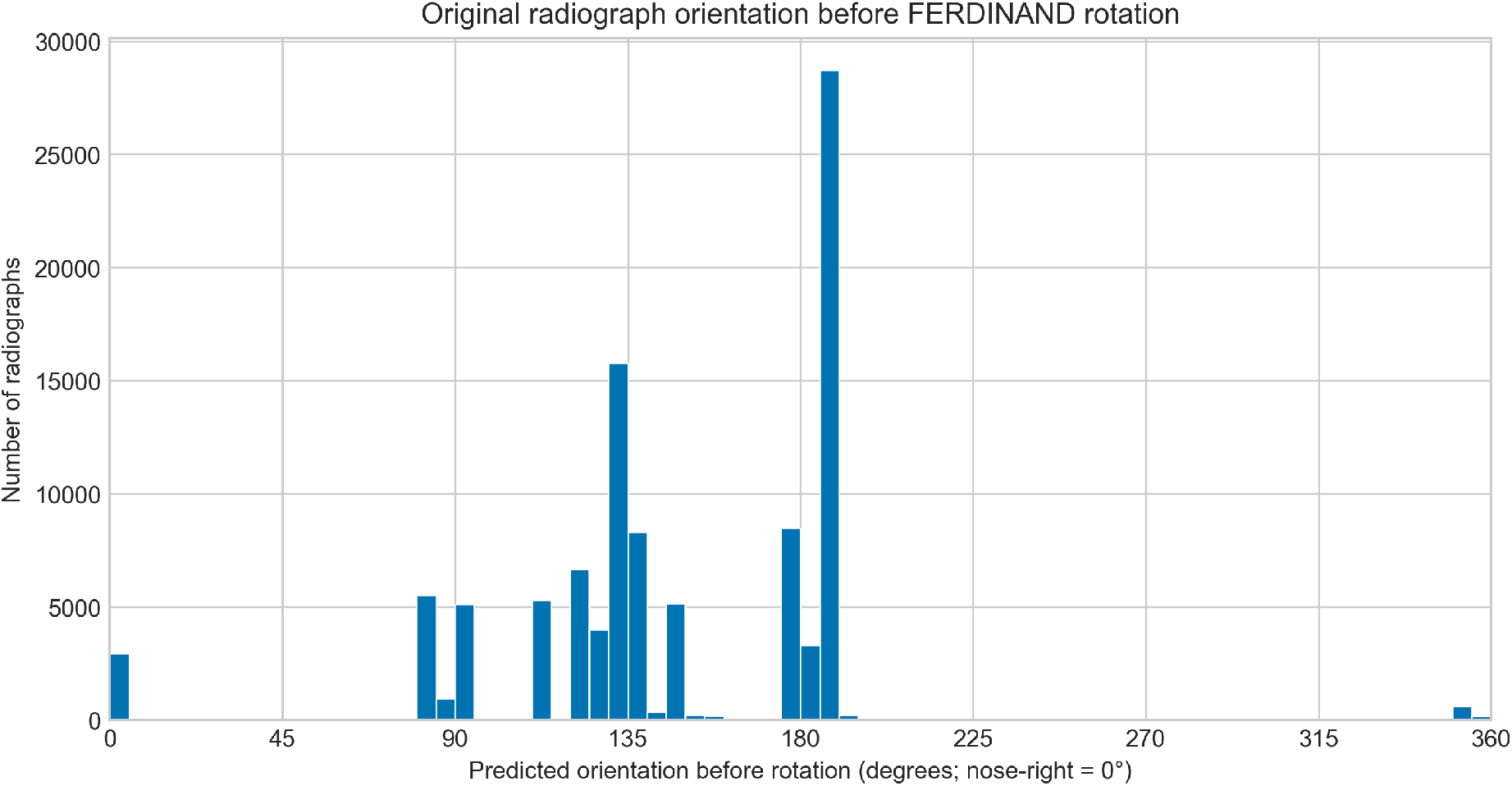
Histogram of original radiograph orientations before FERDINAND rotation, using 5° bins. The nose-right direction is shown at 0°.

The internal structure of the curation problem becomes visible in the embedding space used for semi-automated review. In Figure 4, the JAX example shows that 768D image embeddings form structured centre-level manifolds whose Leiden clusters separate visually distinct image subpopulations. This organisation makes the FERDINAND curation workflow more than a binary filter: clusters can be inspected explicitly in cluster-specific galleries and contact sheets. Representative random samples from JAX further show that lower-numbered clusters capture the expected dorsoventral whole-body radiographs, whereas clusters above 16 increasingly represent off-target image patterns that can be excluded during semi-automated curation. Equivalent centre-specific UMAPs for the remaining centres are provided in Supplementary Figures S1–S9; the full set of JAX cluster contact sheets is provided in Supplementary Figures S10–S13.

**Figure 4:**
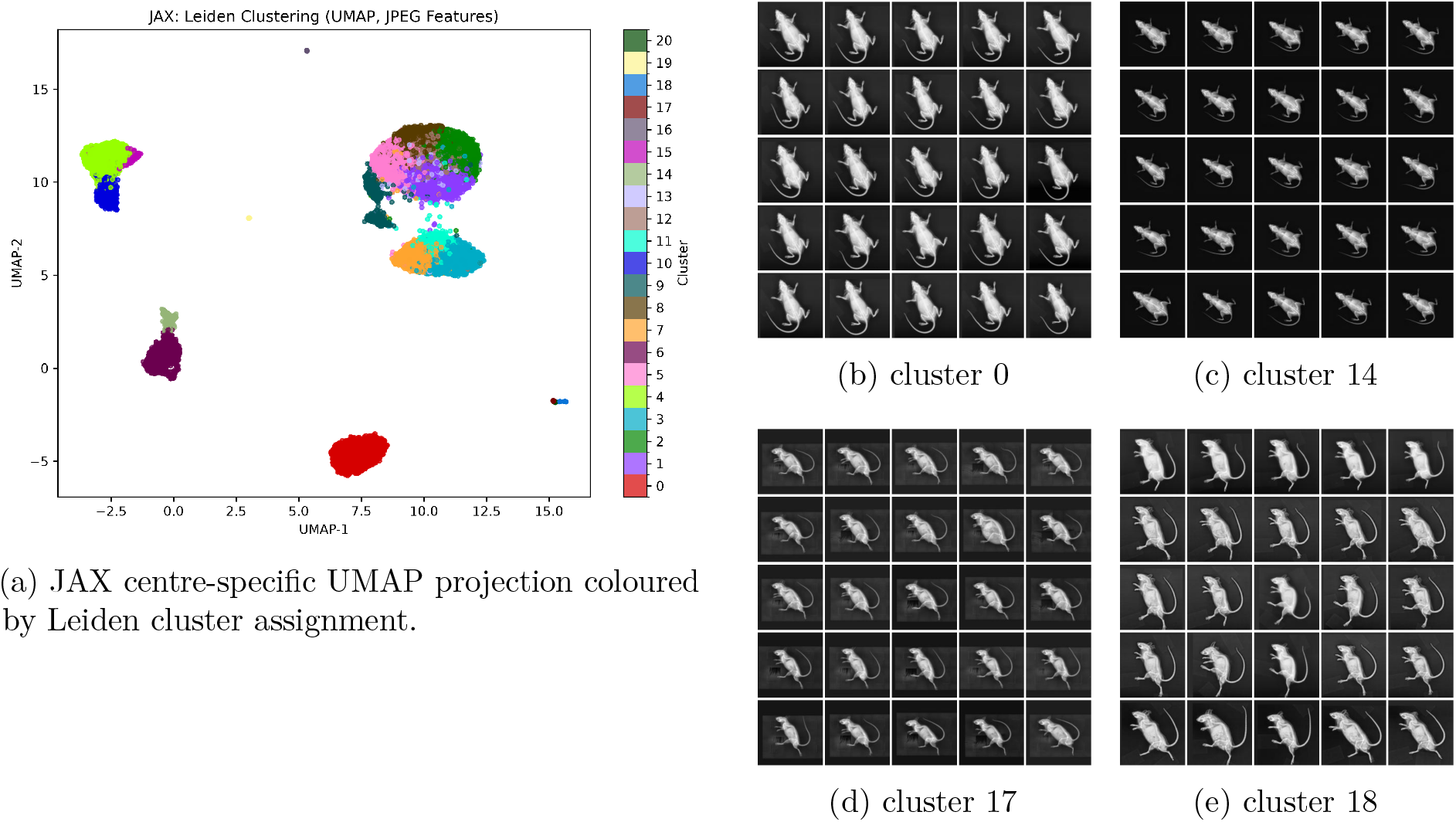
Centre-specific JAX embedding structure and representative random cluster contact sheets used during cluster-guided curation. The left panel (a) shows the JAX UMAP projection based on the 768D FERDINAND image embeddings, with each point representing one radiograph and colours indicating Leiden cluster assignments derived from the centre-specific nearest-neighbour graph. The right panel (b-e) shows random image samples from four JAX Leiden clusters. These contact sheets illustrate that clusters represent distinct visual image patterns and that clusters greater than 16 (d,e) can be excluded because they do not correspond to the expected dorsoventral whole-body radiographs.

At the database level, the current post-curation counts are listed explicitly in Table 6. The complete FERDINAND workflow covers the steps from initial download to final curated output. The practical effect of preprocessing is most readily appreciated at the level of individual images. Figure 5 contrasts original radiographs with their processed counterparts and highlights typical operations such as standardised orientation, contrast enhancement, removal of labels, and suppression of bright metallic markers. These examples complement the aggregate resource summaries by showing how the abstract processing steps represented in Table 5 translate into visually interpretable changes at the level of individual radiographs.

**Figure 5:**
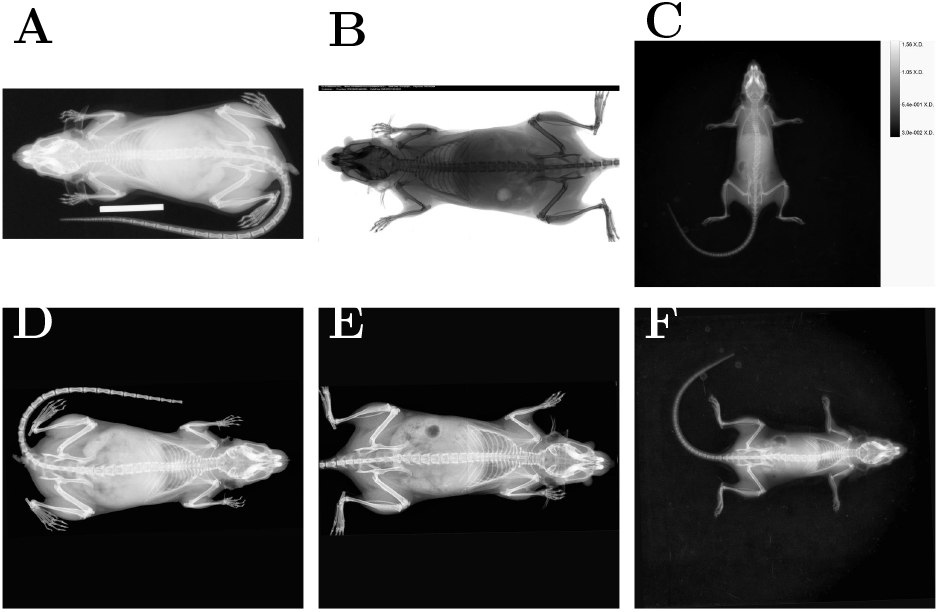
Representative examples of FERDINAND preprocessing. The top row shows original radiographs, and the bottom row the corresponding processed images, so that each column represents one before-versus-after comparison. All processed images were rotated so that the nose points to the right, contrast-enhanced, and padded to square format. Additional steps include **(A,D)** removal of a metallic size marker, **(B,E)** inversion and removal of a top image label, and **(C,F)** removal of a side label.

**Table 6:** Radiograph counts after different steps in the FERDINAND pipeline.

| Stage | Count | % of database |
| --- | --- | --- |
| Radiographs in local FERDINAND database | 104,819 | 100.0 |
| Excluded during cluster-based visual curation | 981 | 0.9 |
| Retained after cluster-based visual curation | 103,837 | 99.1 |
| Overexposed, not rotated | 1,620 | 1.5 |
| Fully processed, rotated | 102,217 | 97.5 |

### 3.2 Proof of Concept: Automated Bone Morphometry

To test whether the harmonised FERDINAND output is suitable for downstream AI-based phenotyping, we applied a separate DeepLabCut-based landmark pipeline for automated bone morphometry. Landmark prediction records are currently available for 102,217 images. For each predicted landmark, image-level *x*- and *y*-coordinates are stored together with a confidence score between 0 and 1. The predicted landmark set covers skull length and width as well as left and right humerus, femur, and tibia landmarks, enabling direct derivation of bone-lengths from FERDINAND-harmonised inputs. Model development compared landmark detection on original radiographs with landmark detection on FERDINAND-processed radiographs using five-fold cross-validation. At the 80% training fraction, FERDINAND preprocessing reduced held-out landmark error and improved mAP, mAR, and threshold-based PCK (Table 7). Figure 6 summarises the cross-validation performance difference between original and FERDINAND-processed radiographs across the principal DeepLabCut error metrics. These end-point gains were modest in the high-performing 80% training setting for threshold-based metrics such as mAP, mAR, and percentage of correct keypoints, but the more consequential effect of FERDINAND preprocessing became apparent in the learning-curve analysis. Specifically, the relative gain from FERDINAND preprocessing was largest when training data were sparse (Table 8). These results indicate that FERDINAND-processed radiographs require less manual annotation to achieve a given level of landmark-detection performance than original radiographs. This data-efficiency effect is visualised directly in Figure 7, where the separation between original and FERDINAND-processed training curves is greatest at reduced training set sizes. Supplementary Figure S14 provide the corresponding threshold-based PCK summaries across training fractions, whereas Figure S15 shows that the advantage was strongest for humerus landmarks.

**Figure 6:**
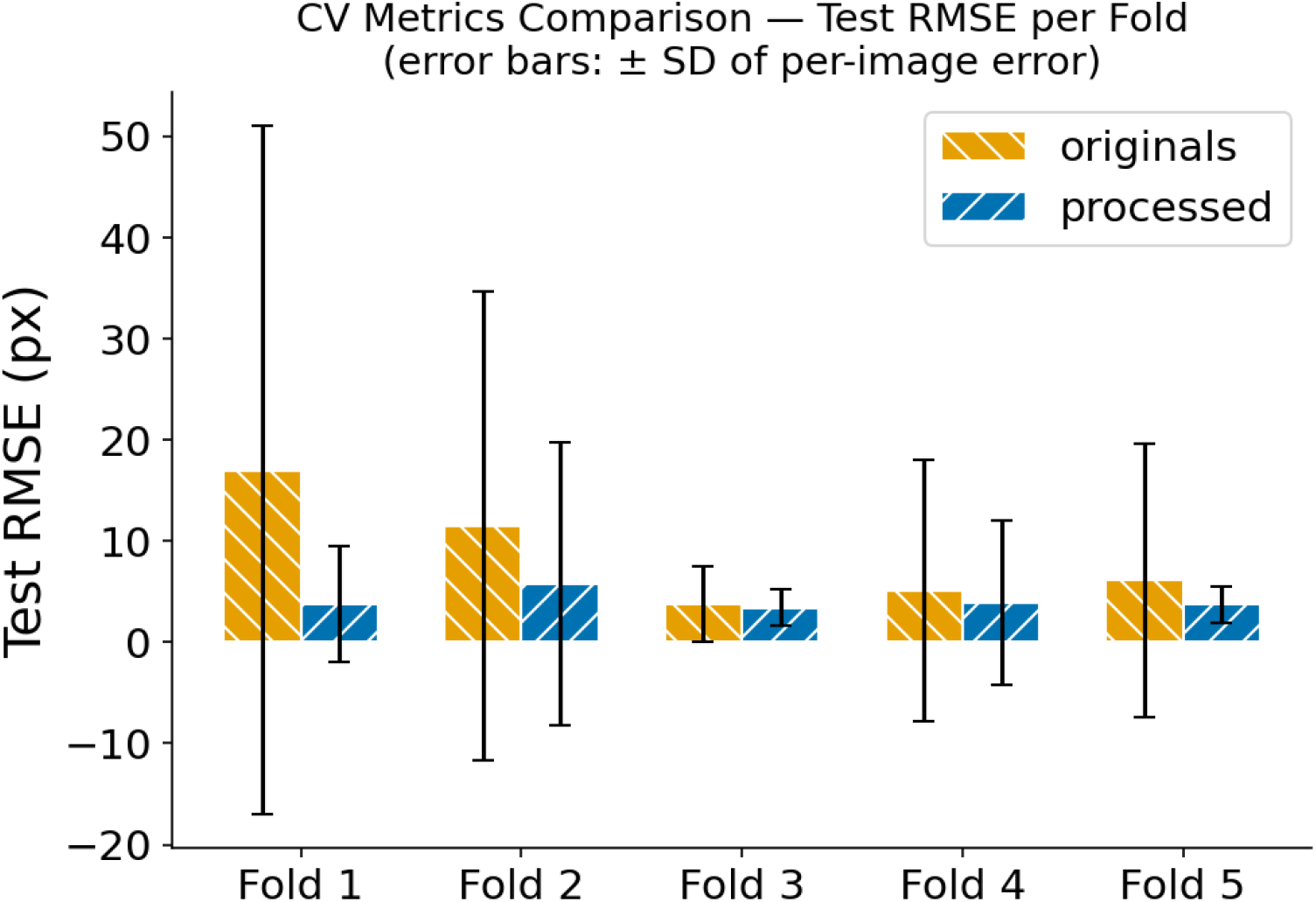
Cross-validation summary of DeepLabCut performance on original and FERDINAND-processed radiographs. Bars show the mean *±* standard deviation across cross-validation folds for the principal test RMSE metrics. FERDINAND preprocessing (preprocessed) lowers test RMSE in every fold and reduces its per-image spread, indicating both more accurate and more consistent downstream DeepLabCut landmark prediction.

**Figure 7:**
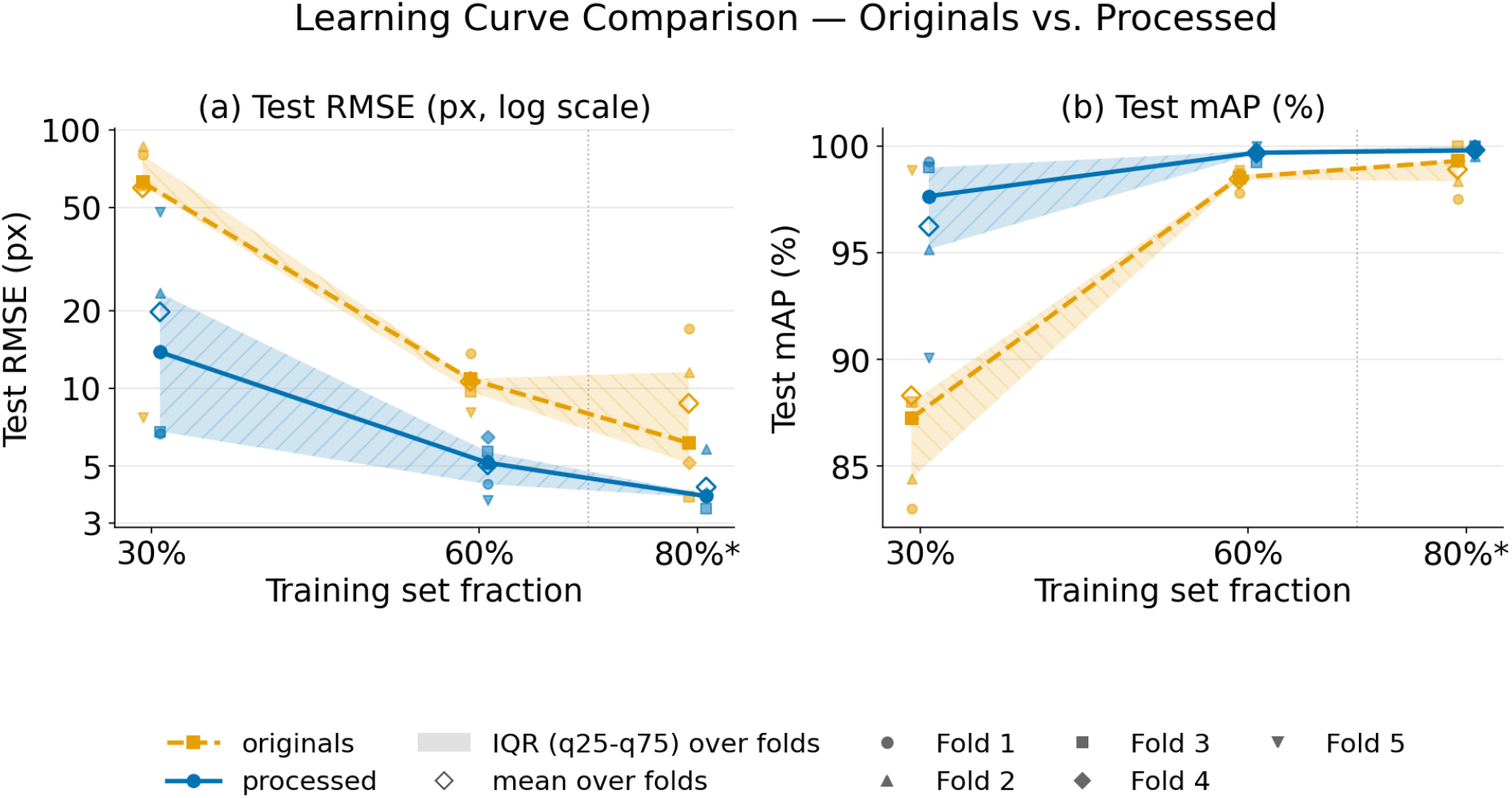
Learning-curve comparison of DeepLabCut performance on original and FERDINAND-processed radiographs, for training set fractions of 30 %, 60 %, and 80 %. (a) Test RMSE (DeepLabCut’s Combined Evaluation RMSE, log scale); (b) Test mAP. Bold lines show the median across the five cross-validation folds; shaded bands show the interquartile range (25th–75th percentile); open diamonds mark the mean; small markers show individual folds. Data points for the two conditions are offset horizontally by ±0.8 percentage points to reduce overlap. The larger separation between both image domains at reduced training set sizes indicates that FERDINAND preprocessing improves data efficiency and reduces the annotation burden required to reach a given level of landmark-detection performance.

**Table 7:** DeepLabCut cross-validation performance at an 80% training fraction. Values for RMSE, mAP, and mAR are mean *±* standard deviation across five folds. PCK values are pooled across 6,113 held-out keypoints. Positive differences favour FERDINAND-processed radiographs, except for RMSE, where a negative difference denotes lower error.

| Metric | Original | FERDINAND-processed | Difference |
| --- | --- | --- | --- |
| Test RMSE (pixels) | $8.72 \pm 5.49$ | $4.13 \pm 0.96$ | $-4.59$ ($-52.6\%$) |
| Test mAP (%) | $98.91 \pm 0.97$ | $99.82 \pm 0.20$ | $+0.91$ |
| Test mAR (%) | $99.03 \pm 0.89$ | $99.86 \pm 0.17$ | $+0.83$ |
| PCK at 5 pixels (%) | 83.8 | 85.9 | $+2.2$ percentage points |
| PCK at 10 pixels (%) | 96.8 | 98.0 | $+1.1$ percentage points |

**Table 8:** DeepLabCut learning-curve results across training fractions. Test RMSE values are mean *±* standard deviation across five cross-validation folds. The relative reduction compares FERDINAND-processed with original radiographs.

| Training fraction | Original RMSE (px) | FERDINAND-processed RMSE (px) | Reduction |
| --- | --- | --- | --- |
| 30% | $59.66 \pm 31.08$ | $19.72 \pm 17.24$ | 67.0% |
| 60% | $10.60 \pm 2.02$ | $5.04 \pm 1.12$ | 52.5% |
| 80% | $8.72 \pm 5.49$ | $4.13 \pm 0.96$ | 52.6% |

For large-scale deployment, the final production model was selected from the FERDINAND-processed variant after a separate quality-control-based comparison of the cross-validation candidates. Deployment configuration, performance, runtime, and coverage are summarised in Table 9. Prediction records were stored in the SQLite `bone_landmarks` table. Of the 103,837 retained radiographs, 102,217 (98.4%) had successfully completed orientation standardisation and received landmark predictions. The remaining 1,620 retained radiographs had not been rotation-corrected due to overexposure. They therefore fell outside the input domain of the production inference workflow. The prediction table contains 102,217 records.

**Table 9:**
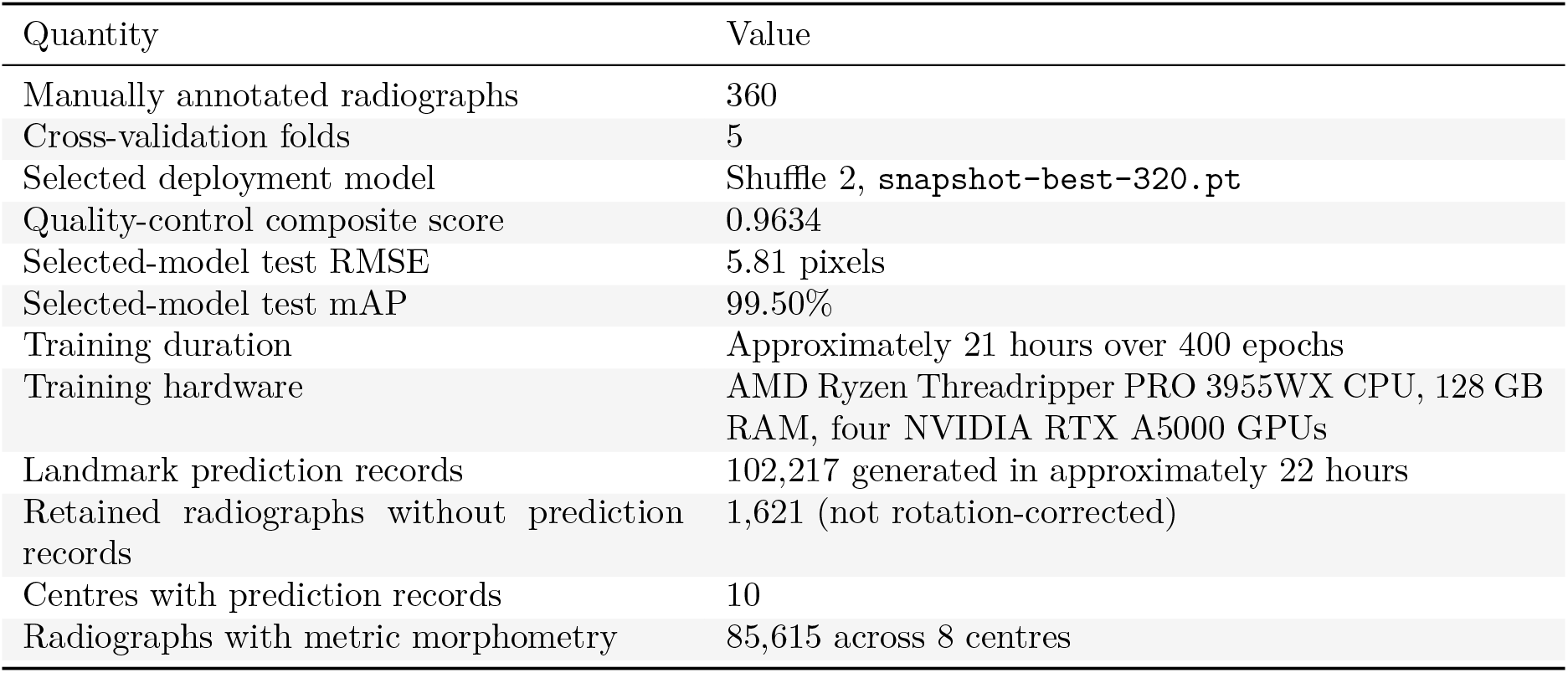
Deployment and coverage of the DeepLabCut-based downstream pipeline. The selected model was chosen after quality-control-based comparison of the cross-validation candidates.

| Quantity | Value |
| --- | --- |
| Manually annotated radiographs | 360 |
| Cross-validation folds | 5 |
| Selected deployment model | Shuffle 2, `snapshot-best-320.pt` |
| Quality-control composite score | 0.9634 |
| Selected-model test RMSE | 5.81 pixels |
| Selected-model test mAP | 99.50% |
| Training duration | Approximately 21 hours over 400 epochs |
| Training hardware | AMD Ryzen Threadripper PRO 3955WX CPU, 128 GB RAM, four NVIDIA RTX A5000 GPUs |
| Landmark prediction records | 102,217 generated in approximately 22 hours |
| Retained radiographs without prediction records | 1,621 (not rotation-corrected) |
| Centres with prediction records | 10 |
| Radiographs with metric morphometry | 85,615 across 8 centres |

For radiographs with metric calibration, derived morphometric measurements are available in millimetres (Table 9). For each retained record, skull length and width as well as left and right humerus, femur, and tibia lengths are available in pixel units and, where metric calibration could be applied, in millimetres. The centre- and sex-stratified humerus distributions shown in Figure 8 demonstrate this downstream utility directly. Within each centre, the female and male distributions remain comparable because they share the same vertical scale, while the grouped layout preserves clear separation between centres and makes the per-stratum sample sizes explicit. The resulting distributions are biologically plausible in several respects. First, left humerus lengths are consistently larger in male mice than in female mice, matching the expected sexual size dimorphism. Second, centre-to-centre shifts in absolute measurements are visible and are compatible with the well-known centre effects [35] documented for numerous other IMPC-derived parameters. Third, the lower panel shows that the signed difference between left and right humerus length remains close to zero across centres and sexes, which is likewise consistent with the expected near-symmetry of paired long bones and with the dedicated bilateral-symmetry quality-control analysis performed on the production predictions. Figure 9 complements these aggregate summaries with a single example radiograph after FERDINAND preprocessing, including the predicted DeepLabCut landmarks. This visual example highlights how the standardised nose-right-oriented FERDINAND output can be used directly for landmark-based quantification on an individual radiograph. As a real-world application example, Figure 10 shows, using the *Bach2* knockout mouse line, how bone lengths derived from FERDINAND-processed IMPC images are consistent with the well-known reduced body size [36] of these mutants. Additional centre- and sex-stratified plots for left femur, left tibia, skull length, skull width, and right-sided long-bone measurements are provided in the Supplementary Figures S16– S22. The resulting distributions show that FERDINAND-preprocessed IMPC radiographs can support large-scale landmark-based phenotyping while preserving centre-level and sex-specific structure for downstream comparative analyses.

**Figure 8:**
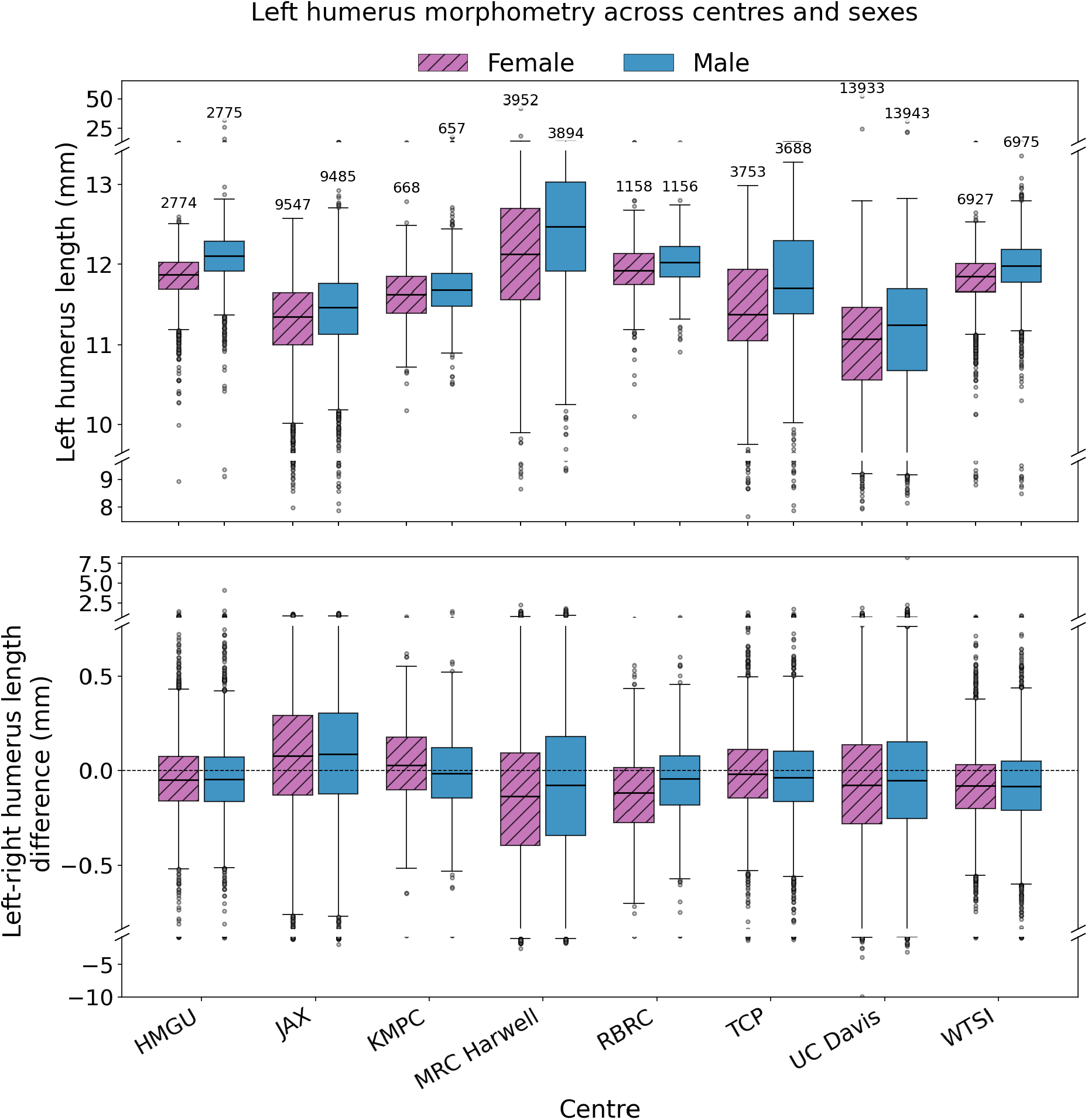
Centre- and sex-stratified humerus morphometry derived from the DeepLabCut-based downstream pipeline. The upper panel shows the distribution of left humerus lengths in millimetres for female and male mice within each centre. The lower panel shows the corresponding signed difference between left and right humerus length for the same centre/sex strata. Numbers above the upper boxplots indicate the number of radiographs contributing to each centre/sex stratum. Pink boxes denote female mice and blue boxes male mice.

**Figure 9:**
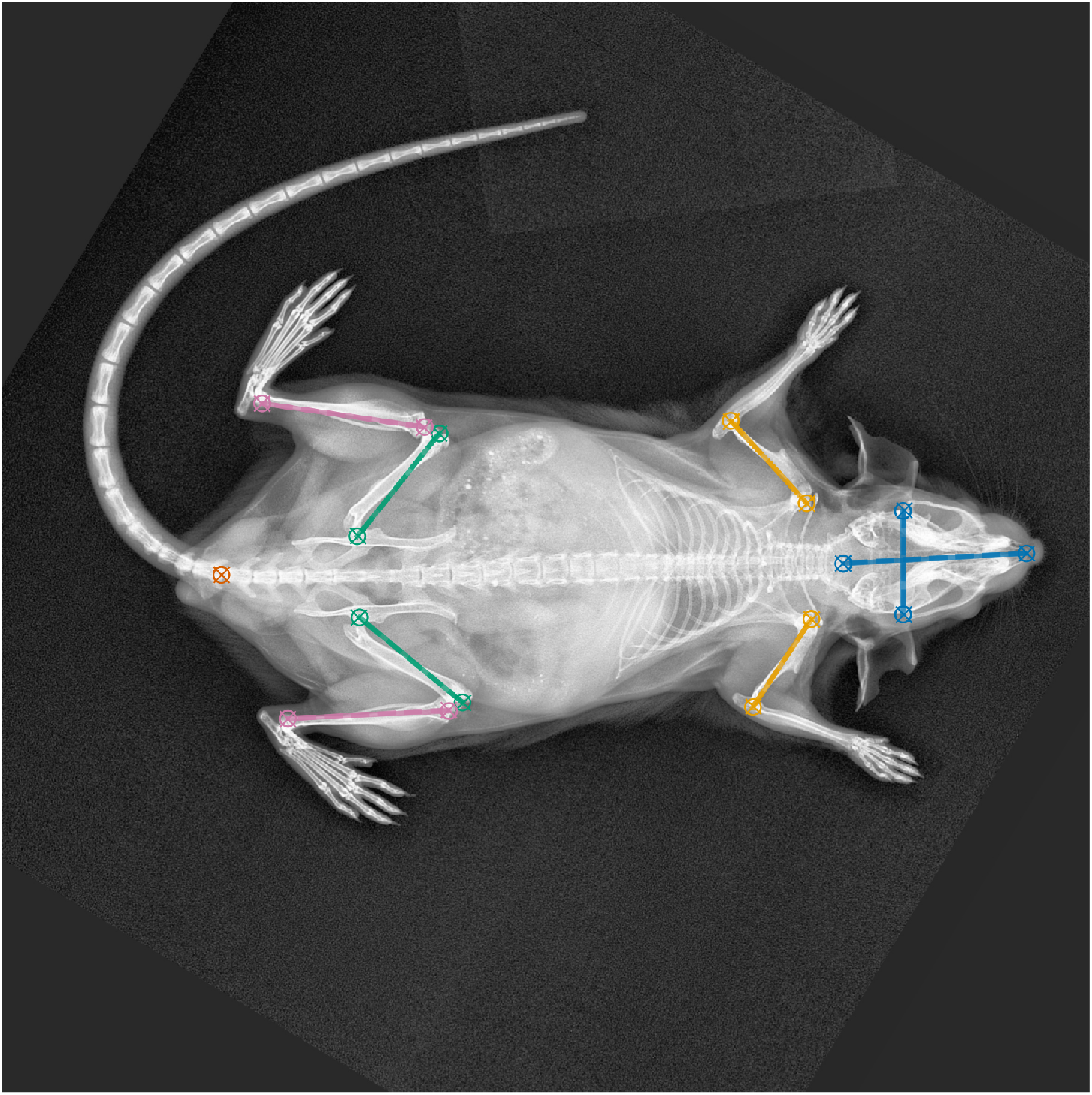
Example of a FERDINAND-processed radiograph with the mouse oriented nose-right. The figure shows DeepLabCut-predicted skeletal landmarks (crosshairs). The centers of the open circles denote the annotated ground truths for comparison. Lengths are calculated as Euclidian distances between corresponding landmarks.

**Figure 10:**
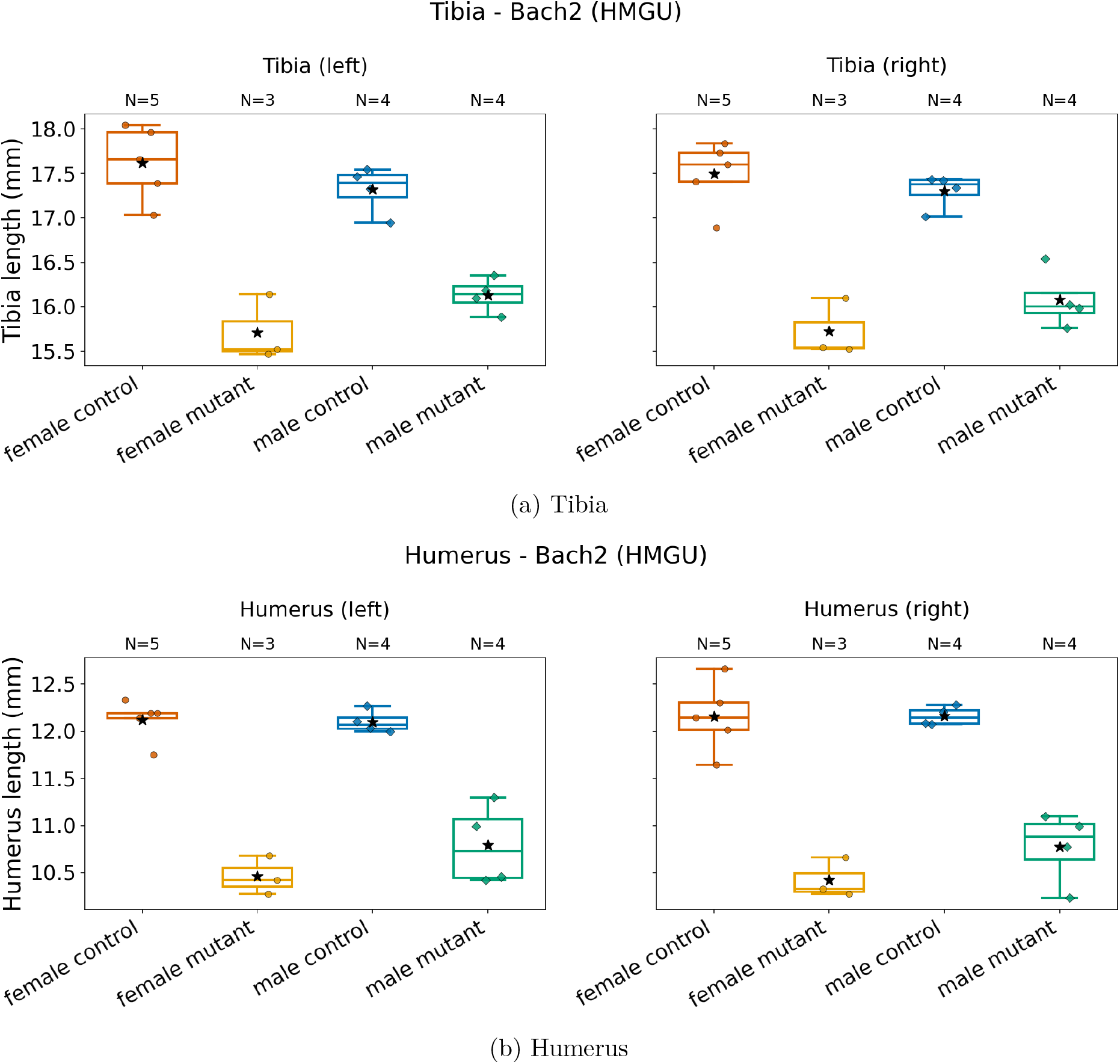
Example of biologically meaningful morphometric separation in the *Bach2* knockout line, measured at HMGU. Length distributions for left and right tibia (top) and humerus (bottom), derived from DeepLabCut landmarks on FERDINAND-processed radiographs, are shown separately for female controls, female mutants, male controls, and male mutants. Across both bones and both sides, *Bach2* −/− mutants show shorter measured lengths than controls, matching the well-established reduced body size phenotype of these mice. Individual data points are shown, and subgroup sizes are given above the plots.

## 4 Discussion

### The harmonised resource

FERDINAND addresses a clear bottleneck in the IMPC radiograph ecosystem: heterogeneous acquisition settings impede direct reuse for machine learning and quantitative image analysis. The framework’s combination of harmonisation, quality control, and curation creates a practical bridge between raw archive content and downstream AI workflows. At the same time, applying the same standardisation across all images also improves human access to the image resource itself: once image orientation, contrast behaviour, and gross technical artefacts are made more uniform, large galleries and contact sheets become substantially easier to inspect. This allows both technical outliers and potentially unusual phenotypes to be identified more efficiently by eye. This contribution is most naturally placed next to the broader literature on harmonisation and quality control of heterogeneous imaging resources. Across medical imaging, scanner-, site-, and protocol-induced variation is widely recognised as a major obstacle for robust multi-centre analysis and machine-learning deployment [37]. Comparable concerns have also motivated technical quality-control methods for radiographs, including automated assessment of positioning, rotation, metadata consistency, and other acquisition defects [38, 39]. FERDINAND extends this logic into the specific setting of IMPC whole-body mouse radiographs, where quality control must be coupled to archive access, metadata harmonisation, centre-aware preprocessing, and manual curation support.

### Position relative to prior quantitative image analysis

Our work is complementary to recent efforts on automated quantitative analysis of murine skeletal radiographs. Rong et al. demonstrated that deep learning can recover bone-length measurements from mouse X-rays at scale [13]. We view this as an important adjacent advance. The distinction is that FERDINAND is not primarily a single-task measurement model; it is an open preprocessing and harmonisation framework intended to make a heterogeneous multi-centre image resource accessible for a wider range of downstream methods. Likewise, the broader landmark-detection literature in radiography [14] shows that supervised localisation of anatomical points is technically mature in several domains, but it generally assumes more standardised inputs than those encountered in the IMPC radiograph archive. In the present study, the downstream DeepLabCut-based bone morphometry serves as an application example showing that FERDINAND can provide adequately harmonised input for a successful AI measurement workflow. In the current project state, this downstream comparison is informative not only because landmark-based morphometry becomes feasible at scale, but also because it clarifies where FERDINAND’s strongest downstream benefit lies. In the present DeepLabCut analysis, the gain in final high-level detection metrics in the fully trained setting was comparatively modest, whereas the improvement in data efficiency was much more informative: FERDINAND-processed images supported substantially better performance at reduced training-set sizes. This implies that fewer expert annotations are needed to reach a given level of landmark quality than when training on original images. That point is practically important because expert landmark annotation is costly, and any reduction in annotation burden substantially improves the accessibility of quantitative phenotyping workflows. A dedicated manuscript on that morphometry pipeline is in preparation.

### The publication workflow

An additional strength of the present work is its emphasis on reproducibility. The analytical workflow is organised so that quantitative summaries, tables and figures can be regenerated in a controlled manner from the underlying data and processing pipeline.

### Current limitations

Several scientific limitations should be considered. FERDINAND harmonises and curates heterogeneous IMPC radiographs, but it does not eliminate the inherent constraints of 2D projection imaging. In particular, bone lengths derived from 2D radiographs may be influenced by projection-dependent shortening, so such measurements should be regarded as imaging-based proxies rather than exact anatomical ground truth. Even so, when images are produced, processed, and analysed under the same conditions, these proxy measures can still yield biologically interpretable relative differences, as shown here in the *Bach2 −/−* morphometry example. This application should be seen as one downstream use case rather than the defining purpose of FERDINAND: its broader value lies in converting a visually and technically heterogeneous archive into a more uniform, AI-ready resource that is easier to analyse both computationally and by human inspection. The DeepLabCut comparison further suggests that FERDINAND’s main contribution is not necessarily to transform an already strong landmark model into a dramatically more accurate one, but to reduce nuisance variation in a way that makes downstream learning more data-efficient. The processed images may therefore also support future analyses of soft tissues, body composition, skeletal shape, and other radiograph-derived phenotypes. Absolute conversion to millimetres remains limited by the availability of resolution metadata, and centre-specific preprocessing choices may need revision if future data sources differ substantially.

### Outlook

The harmonised resource provides a basis for future work on broader morphometric descriptors, automated anomaly screening, and additional AI-supported phenotyping tasks, potentially including radiomics-style analyses, beyond the initial proof of concept. Although the present downstream demonstration uses a DeepLabCut-based landmark pipeline, we expect the same improvements in image quality and cross-centre standardisation to be beneficial for other downstream analysis strategies as well, including methods that rely on visual embeddings, shape analysis, segmentation, or other learned image representations.

## ACKNOWLEDGEMENTS

We thank the International Mouse Phenotyping Consortium for making the data resource publicly available. We also thank Dr. Adrián Sanz-Moreno from the German Mouse Clinic for providing terminal bone landmarks for DeepLabCut training.

## DECLARATIONS

## CRediT authorship contribution statement

Conceptualization (RS, HM, MHA, ES); Data curation (RS, ES); Formal analysis (RS, HM, ES); Funding acquisition (MHA); Investigation (HM, ES); Methodology (RS, HM, MHA, ES); Project administration (HM, MHA, ES); Resources (MHA, ES); Software (RS, HM, ES); Supervision (HM, MHA, ES); Validation (RS, HM, ES); Visualization (RS, HM, ES); Writing – original draft (RS, HM, ES); Writing – review & editing (RS, HM, MHA, ES).

## Declaration of Generative AI Use

The authors used AI-assisted coding agents (OpenAI Codex and Anthropic Claude Code) in a human-in-the-loop workflow to support Python code generation during development of the analysis pipeline. Large language models (ChatGPT, Claude) were used for language editing of the manuscript. As the project spanned over one year, multiple model versions were employed over time. AI tools were not used to generate the underlying data, perform data analysis autonomously, or make scientific interpretations or conclusions. All AI-generated code was manually reviewed, tested, and validated by the authors. All AI-assisted text edits were critically assessed and revised. The authors retain full responsibility for the integrity and accuracy of the work.

## Data and Code Availability

The FERDINAND source code is available at https://github.com/ExperimentalGenetics/FERDINAND.

## Animal Ethics

No animal experiments were performed as part of this study. This work analysed radiographic and metadata resources generated and released by the International Mouse Phenotyping Consortium.

