## Supplementary Material for "FERDINAND: An Open Framework for Download and AI-ready Preprocessing of IMPC 2D Radiographs"

Table S1: **Full list of distinct original image geometries represented in the current FERDINAND resource**.

| Width | Height | Count |
|---|---|---|
| 2,368 | 2,340 | 47,083 |
| 3,072 | 2,048 | 38,306 |
| 2,048 | 1,024 | 7,173 |
| 2,364 | 2,048 | 3,694 |
| 2,048 | 2,048 | 2,702 |
| 2,048 | 1,064 | 2,629 |
| 3,072 | 2,148 | 1,303 |
| 2,500 | 2,048 | 1,100 |
| 2,948 | 2,048 | 285 |
| 5,888 | 4,608 | 260 |
| 2,324 | 2,048 | 192 |
| 2,048 | 1,054 | 30 |
| 2,800 | 2,048 | 20 |
| 384 | 256 | 7 |
| 2,516 | 2,048 | 6 |
| 2,612 | 2,048 | 6 |
| 2,408 | 2,048 | 5 |
| 2,208 | 2,048 | 4 |
| 2,428 | 2,048 | 3 |
| 2,356 | 2,048 | 2 |
| 2,380 | 2,048 | 2 |
| 2,416 | 2,048 | 2 |
| 2,228 | 2,048 | 1 |
| 2,308 | 2,048 | 1 |
| 2,340 | 2,048 | 1 |
| 2,424 | 2,048 | 1 |
| 3,144 | 2,482 | 1 |

### Supplementary FERDINAND Curation Figures

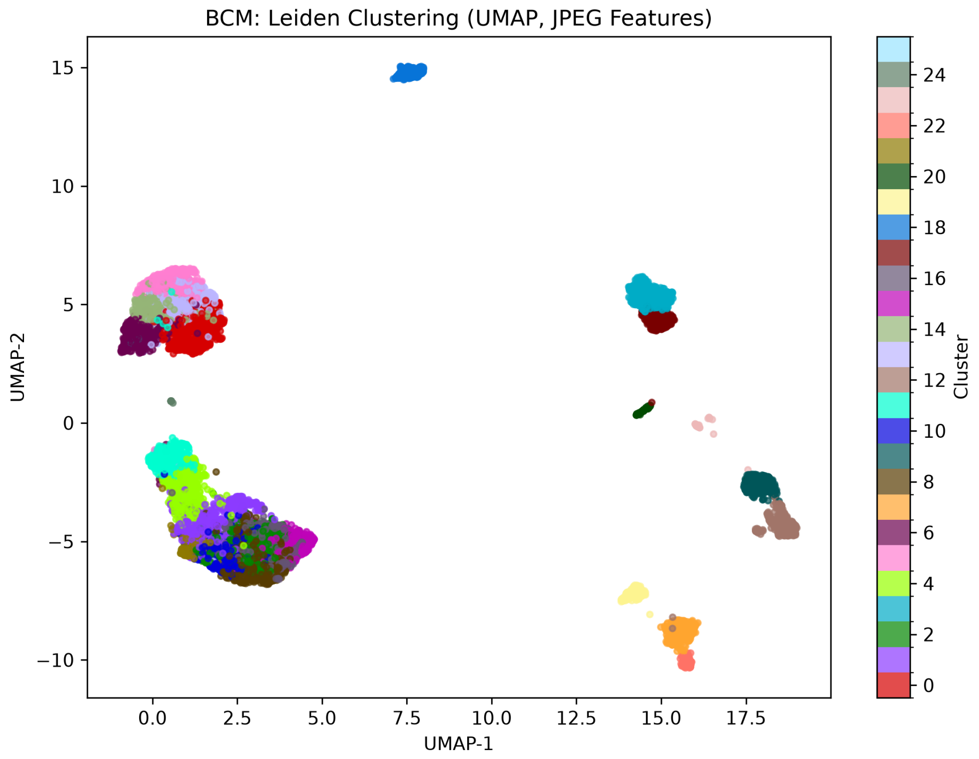


Figure S1: **Supplementary centre-specific UMAP projection for BCM based on the FERDINAND curation embeddings**. Colours indicate Leiden cluster assignments.

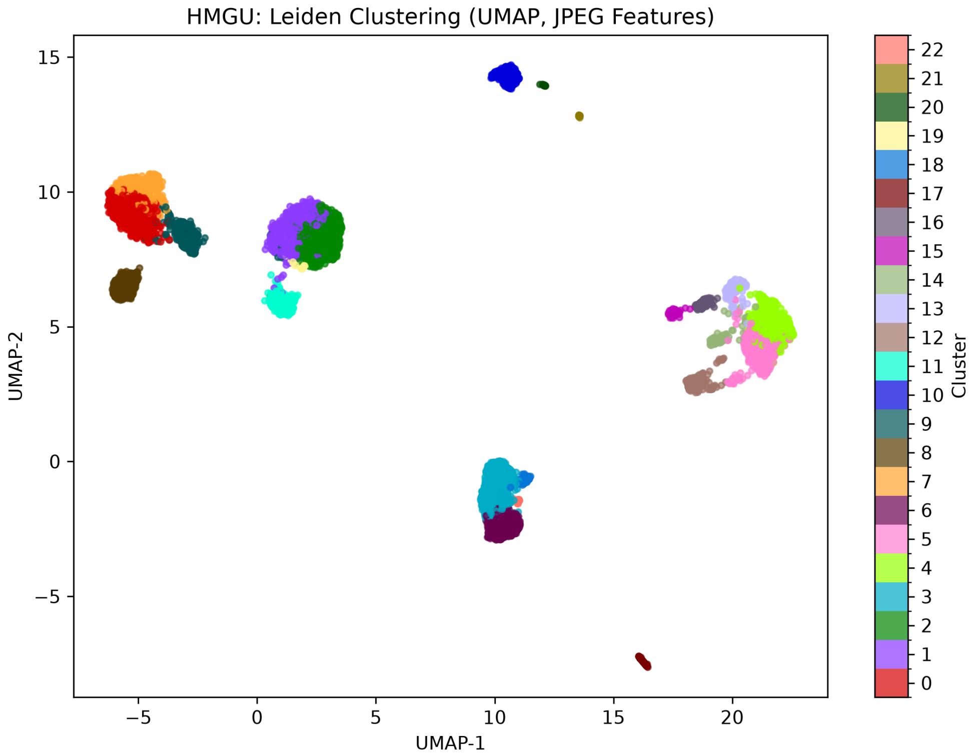


Figure S2: **Supplementary centre-specific UMAP projection for HMGU based on the FERDINAND curation embeddings**. Colours indicate Leiden cluster assignments.

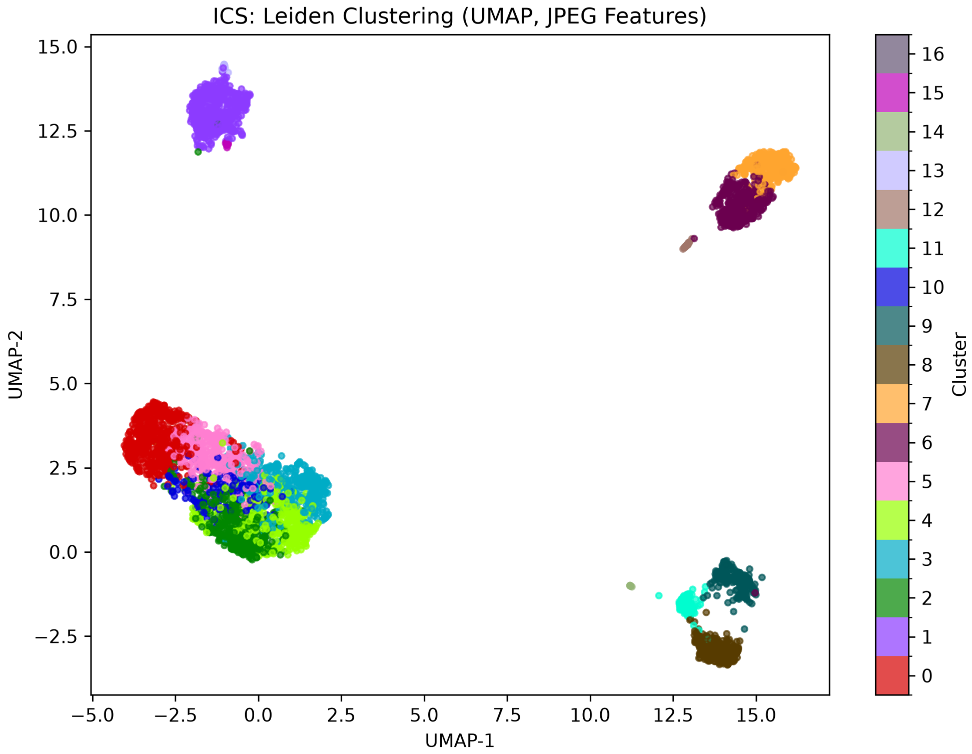


Figure S3: **Supplementary centre-specific UMAP projection for ICS based on the FERDINAND curation embeddings**. Colours indicate Leiden cluster assignments.

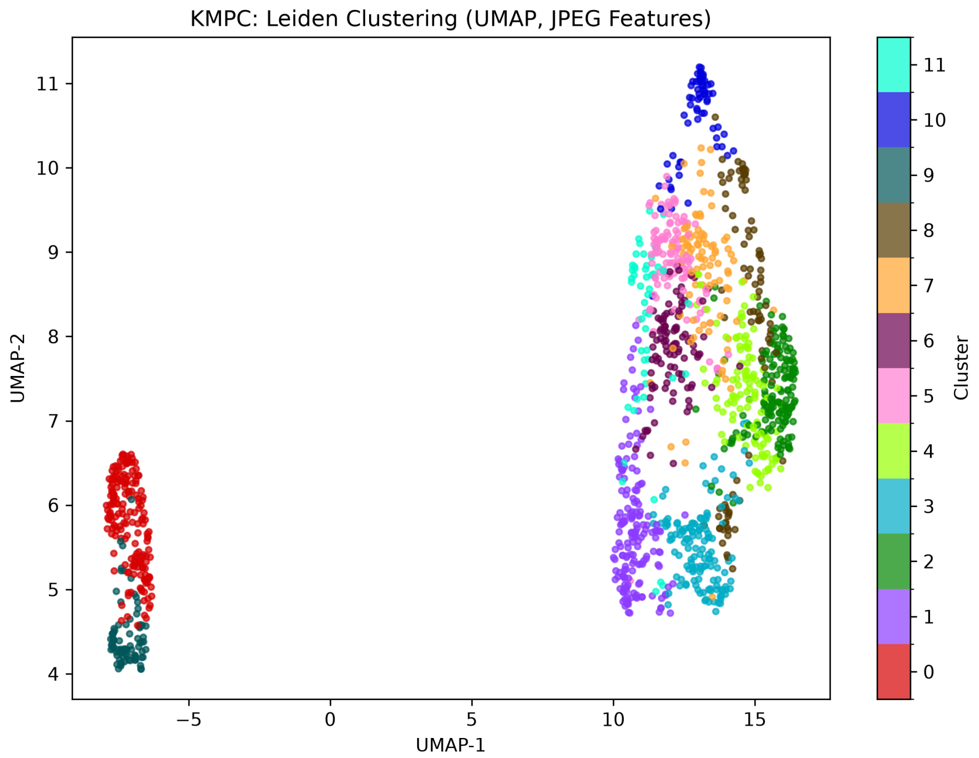


Figure S4: **Supplementary centre-specific UMAP projection for KMPC based on the FERDINAND curation embeddings**. Colours indicate Leiden cluster assignments.

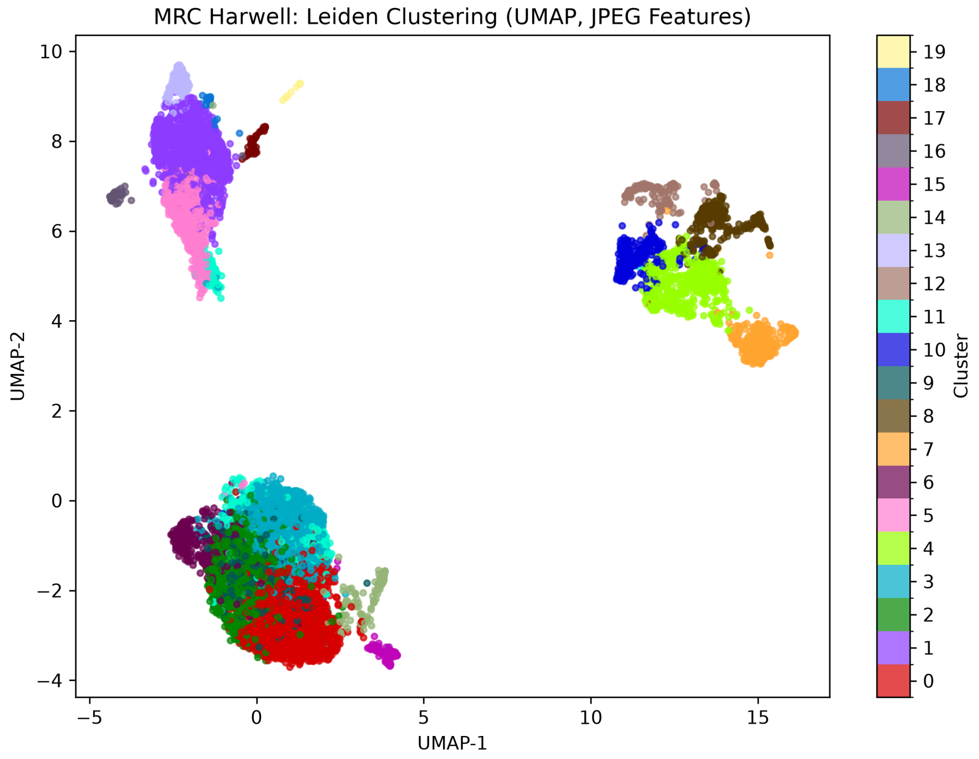


Figure S5: **Supplementary centre-specific UMAP projection for MRC Harwell based on the FERDINAND curation embeddings**. Colours indicate Leiden cluster assignments.

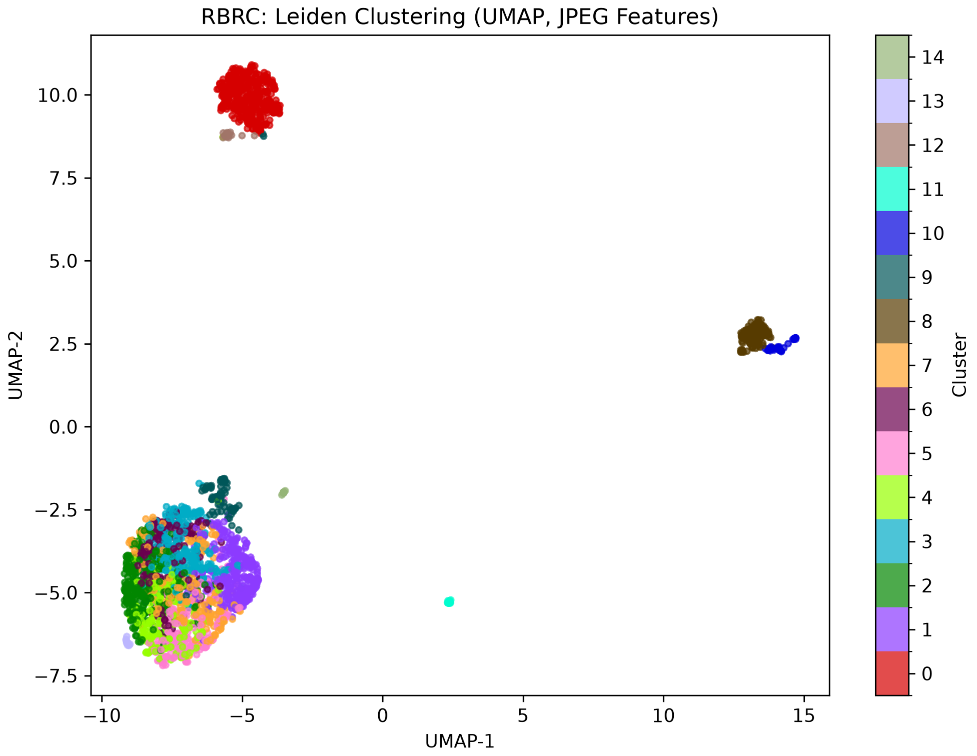


Figure S6: **Supplementary centre-specific UMAP projection for RBRC based on the FERDINAND curation embeddings**. Colours indicate Leiden cluster assignments.

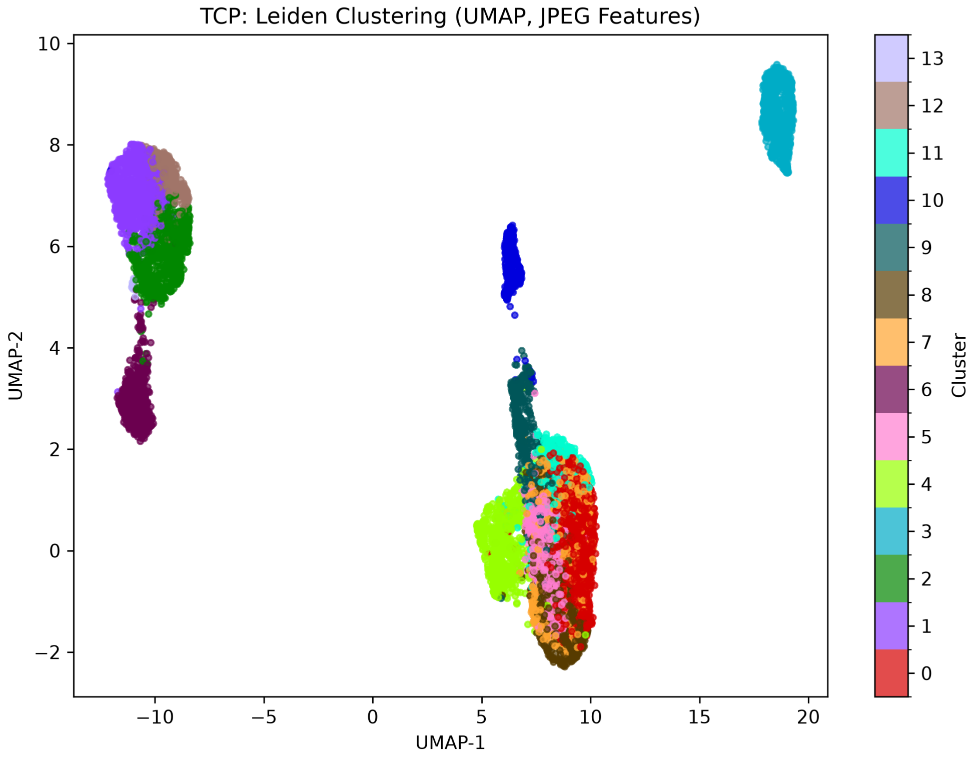


Figure S7: **Supplementary centre-specific UMAP projection for TCP based on the FERDINAND curation embeddings**. Colours indicate Leiden cluster assignments.

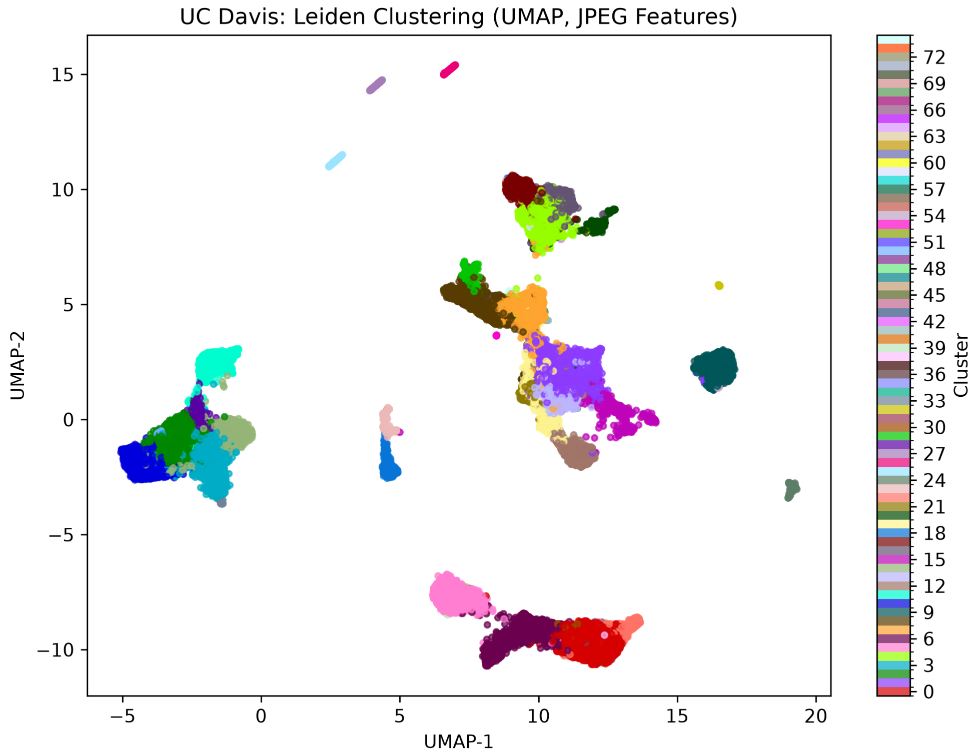


Figure S8: **Supplementary centre-specific UMAP projection for UC Davis based on the FERDINAND curation embeddings**. Colours indicate Leiden cluster assignments.

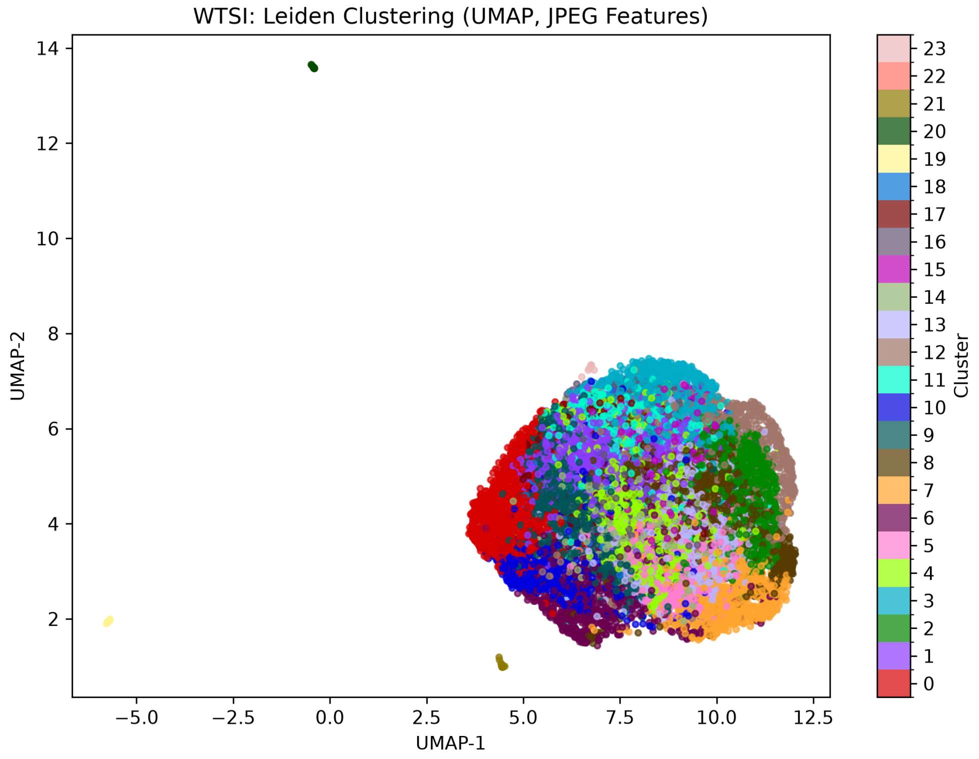


Figure S9: **Supplementary centre-specific UMAP projection for WTSI based on the FERDINAND curation embeddings**. Colours indicate Leiden cluster assignments.

#### Supplementary JAX Cluster Contact Sheets

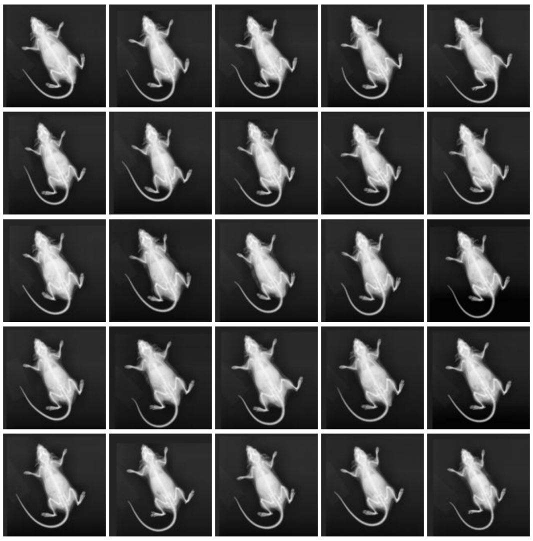

(a) JAX - cluster 0

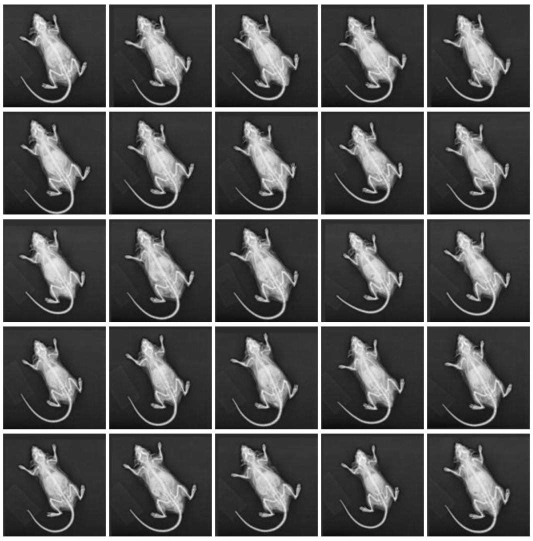

(b) JAX - cluster 1

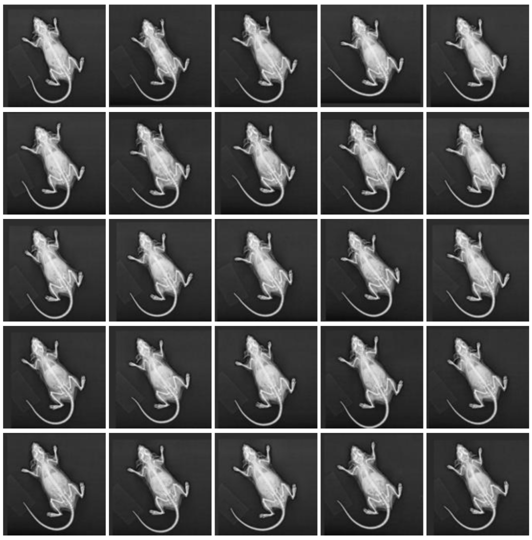

(c) JAX - cluster 2

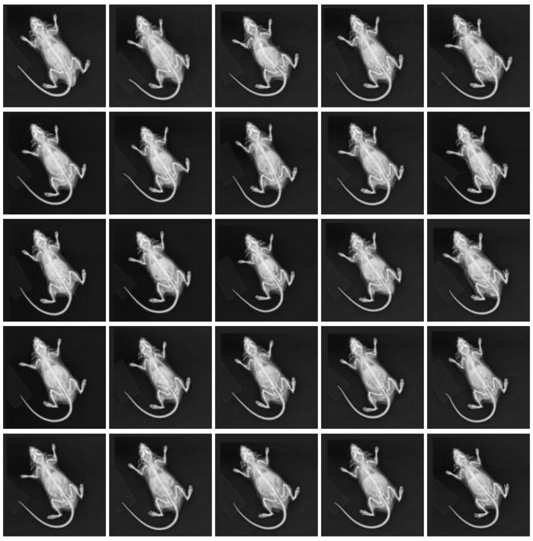

(d) JAX - cluster 3

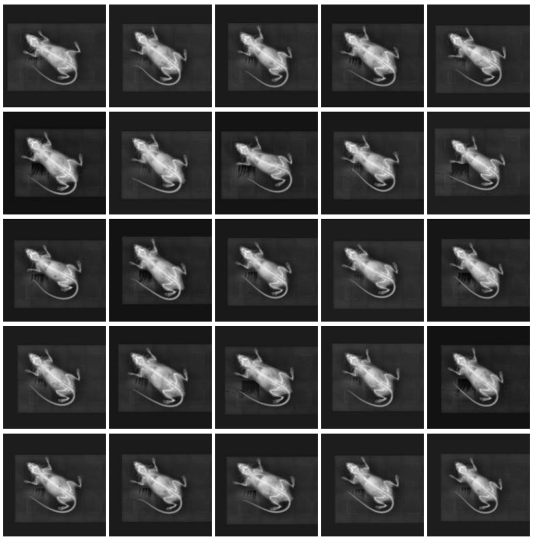

(e) JAX - cluster 4

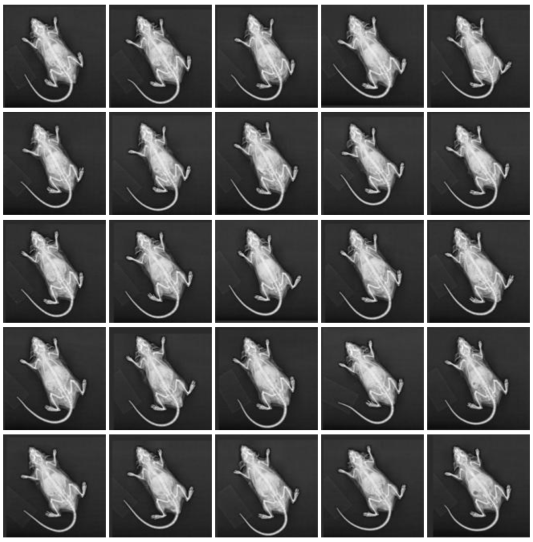

(f) JAX - cluster 5

Figure S10: Exemplary random image samples from the Leiden clusters of the JAX image feature vectors. The clusters represent distinct image patterns within the centre-specific embedding manifold.

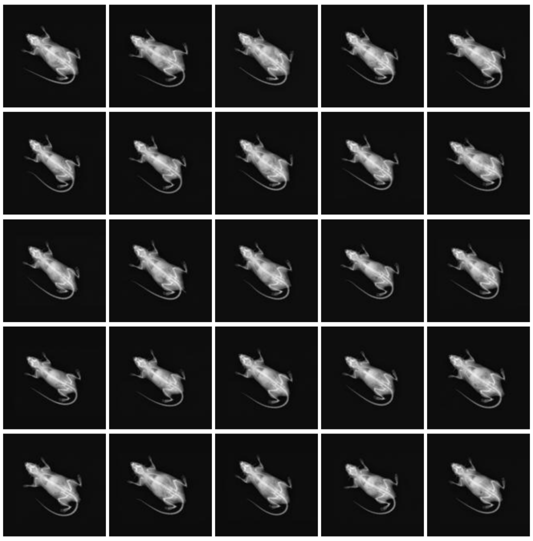

(a) JAX - cluster 6

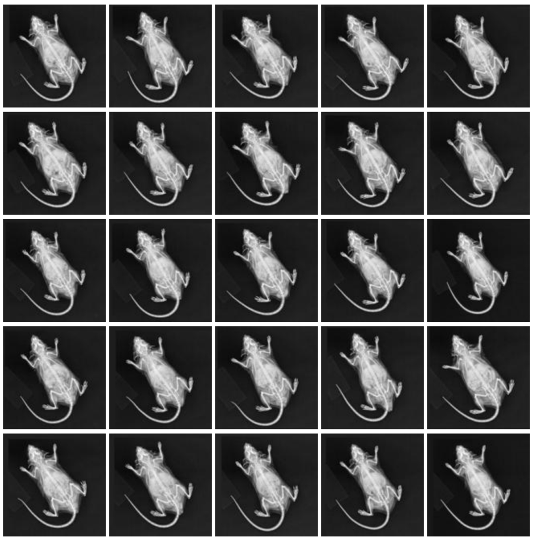

(b) JAX - cluster 7

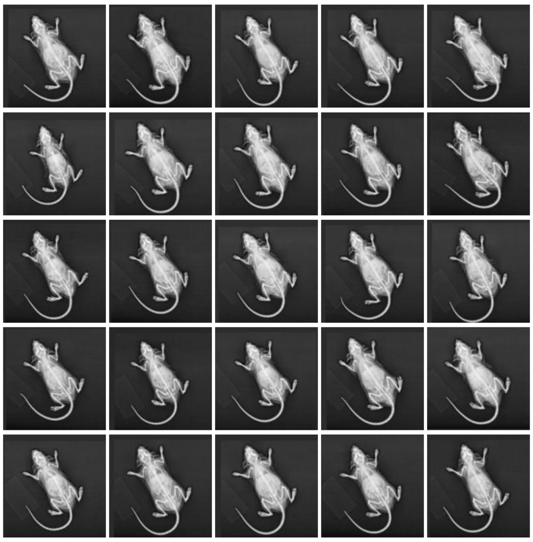

(c) JAX - cluster 8

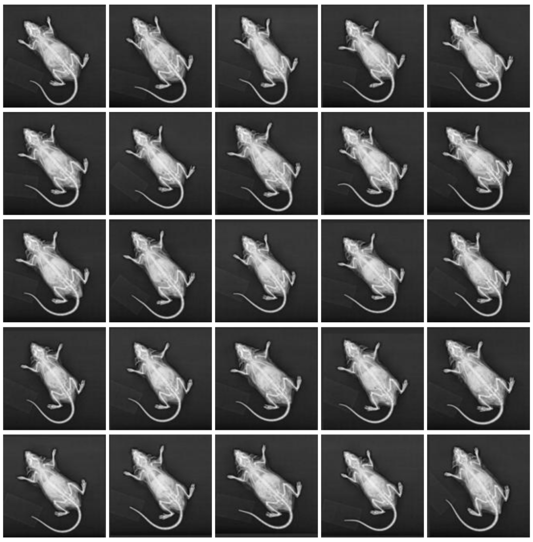

(d) JAX - cluster 9

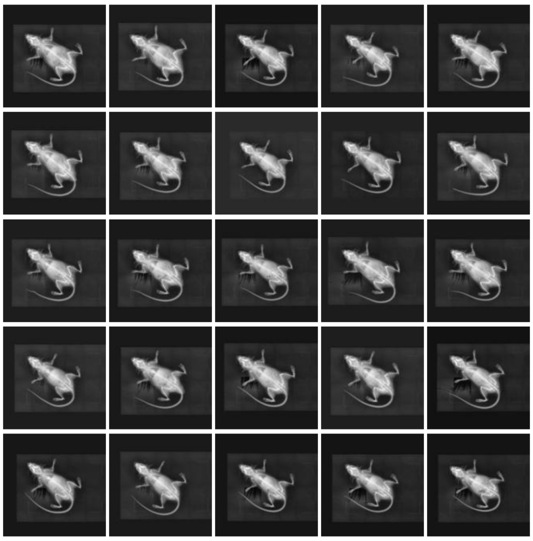

(e) JAX - cluster 10

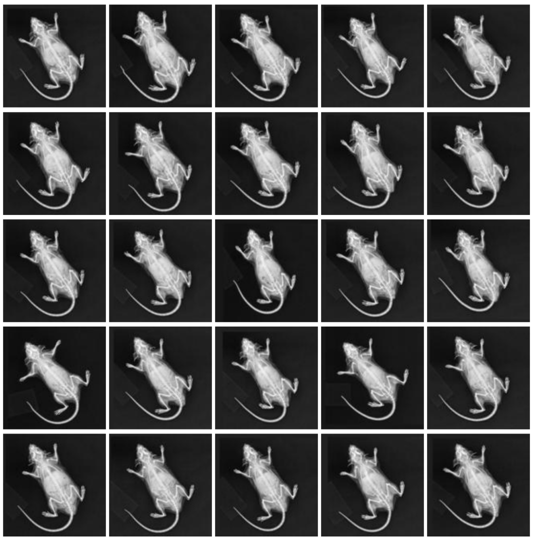

(f) JAX - cluster 11

Figure S11: Exemplary random image samples from the Leiden clusters of the JAX image feature vectors (continued).

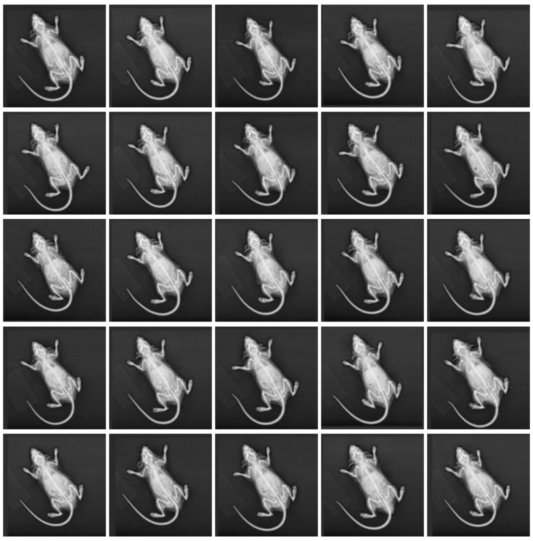

(a) JAX - cluster 12

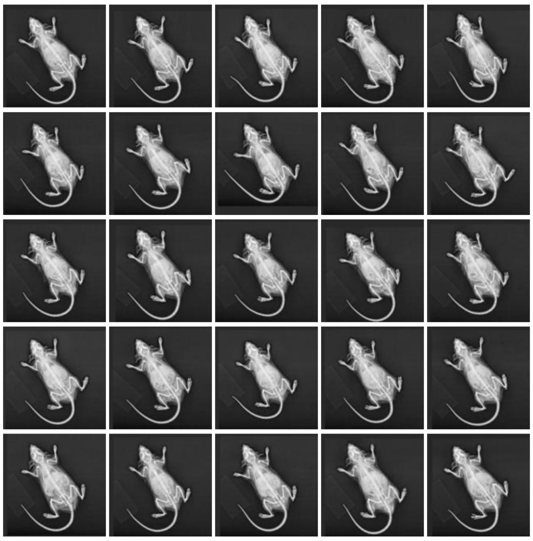

(b) JAX - cluster 13

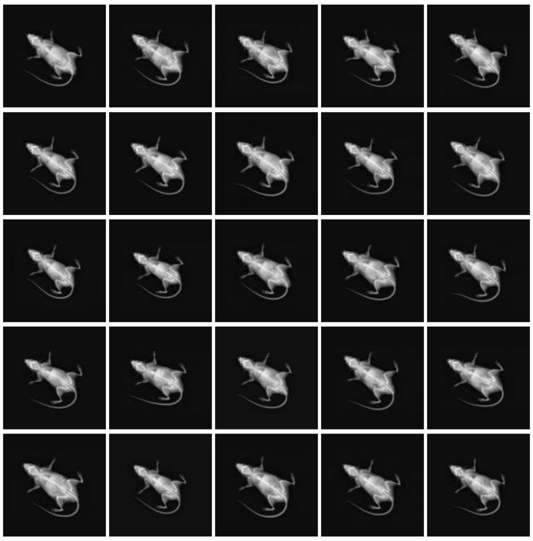

(c) JAX - cluster 14

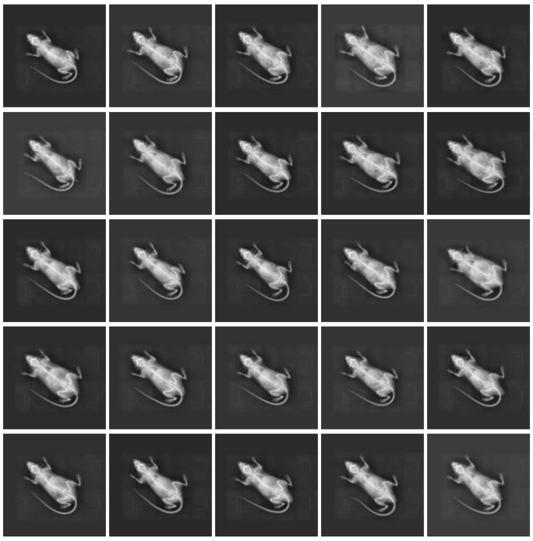

(d) JAX - cluster 15

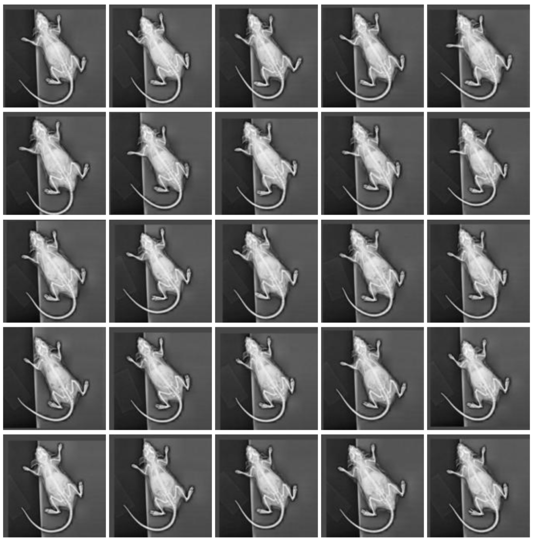

(e) JAX - cluster 16

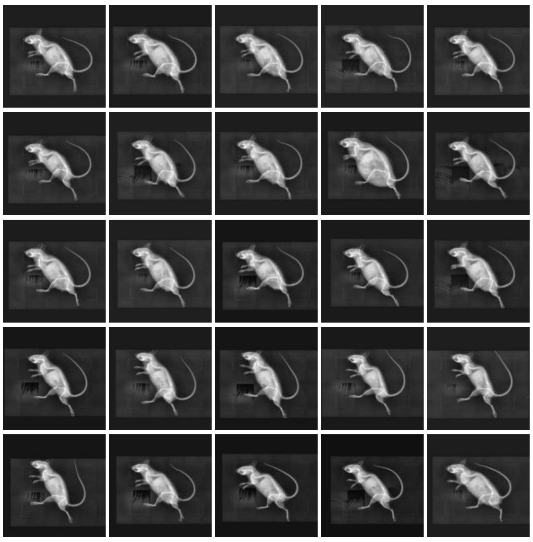

(f) JAX - cluster 17

Figure S12: **Exemplary random image samples from the Leiden clusters of the JAX image feature vectors (continued)**. Images from cluster 17 were excluded because they show lateral instead of the expected dorsoventral whole-body radiographs.

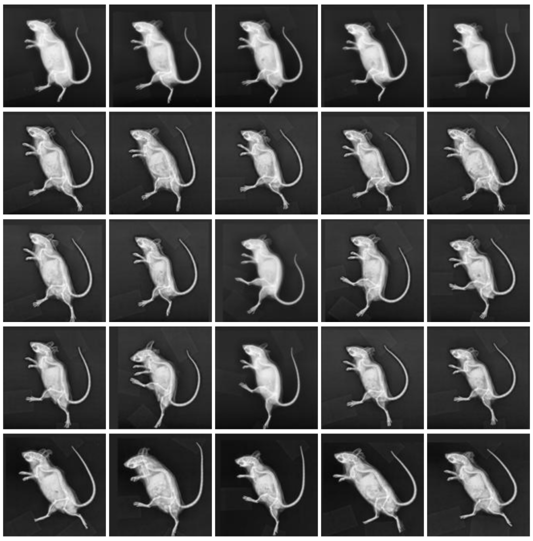

(a) JAX - cluster 18

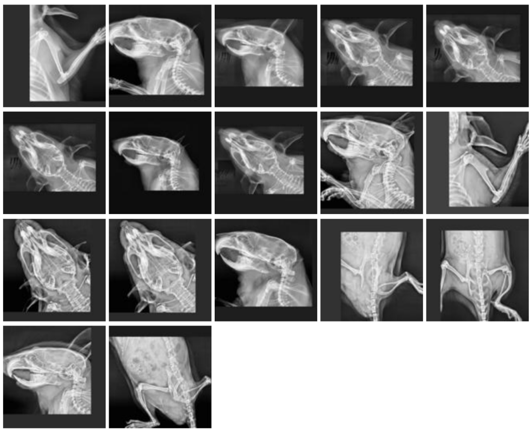

(b) JAX - cluster 19

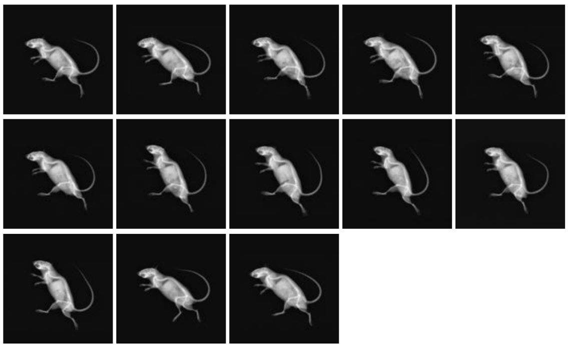

(c) JAX - cluster 20

Figure S13: **Exemplary random image samples from the Leiden clusters of the JAX image feature vectors (continued)**. Images from clusters 18, 19, and 20 were excluded. Cluster 19 and 20 show lateral whole-body images, cluster 18 only shows partial images.

#### Supplementary DeepLabCut Validation Figures

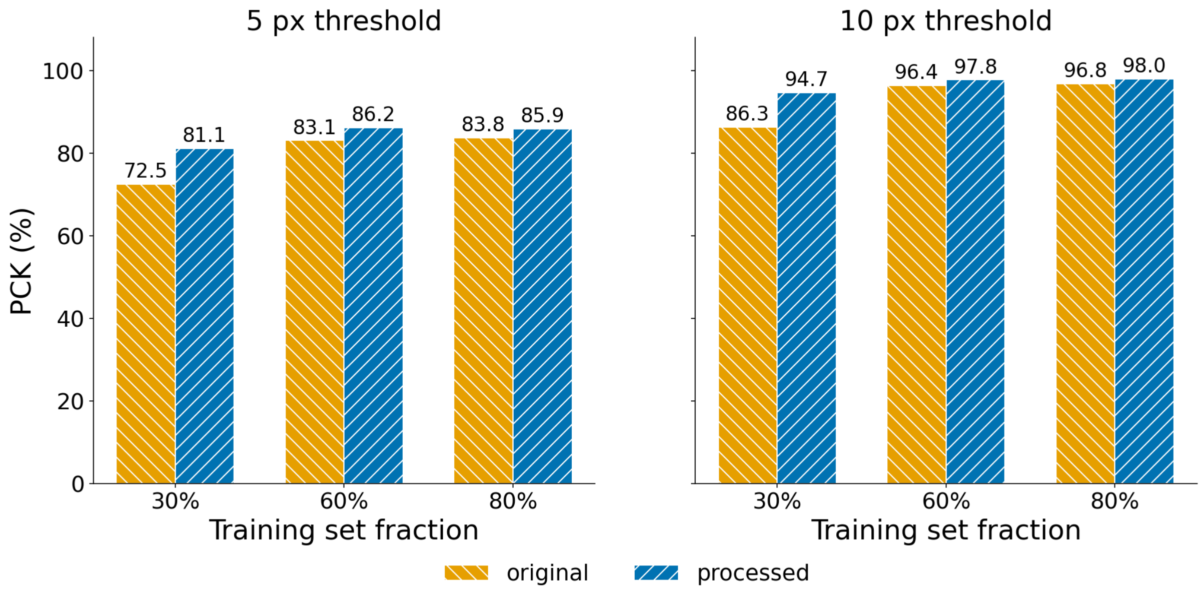


Figure S14: **Percentage of Correct Keypoints (PCK) for DeepLabCut models trained on original and FERDINAND-processed radiographs across training fractions of 30%, 60%, and 80%**. PCK values are pooled over all keypoints and all test images of the five cross-validation folds. The advantage of FERDINAND preprocessing is present for both the 5-pixel and 10-pixel correctness thresholds and is most pronounced at reduced training set size.

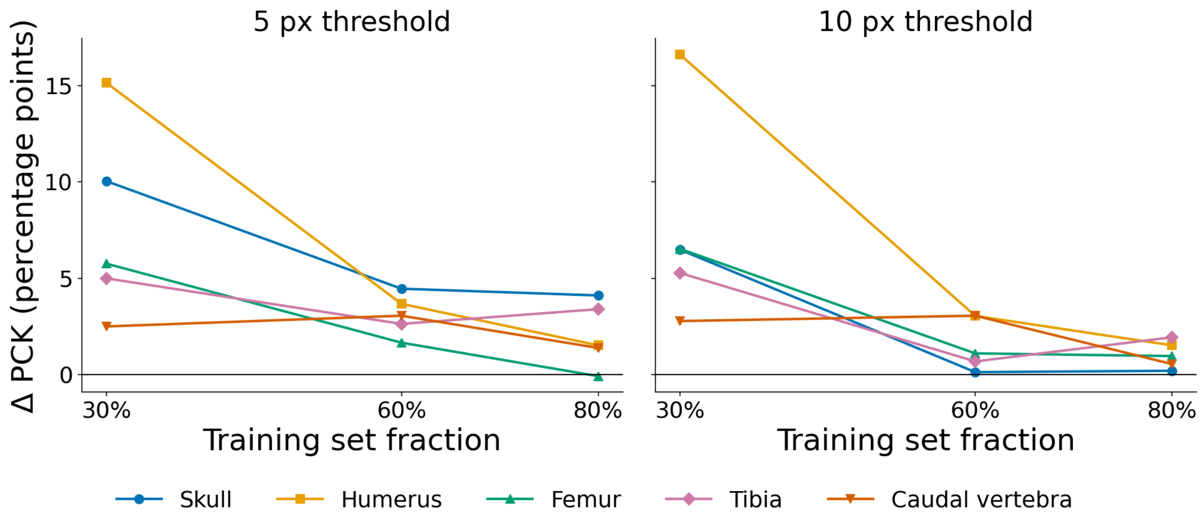


Figure S15: **Difference in Percentage of Correct Keypoints (Δ PCK, processed minus originals) per anatomical group (skull, humerus, femur, tibia, caudal vertebra) across training-set fractions of 30%, 60%, and 80%, shown separately for the 5-pixel and 10-pixel correctness thresholds**. PCK values are pooled over all test keypoints of the five cross-validation folds. Positive values indicate that FERDINAND preprocessing improves keypoint accuracy for that group and training fraction; the horizontal line at zero marks no difference.

### Supplementary Morphometry Figures

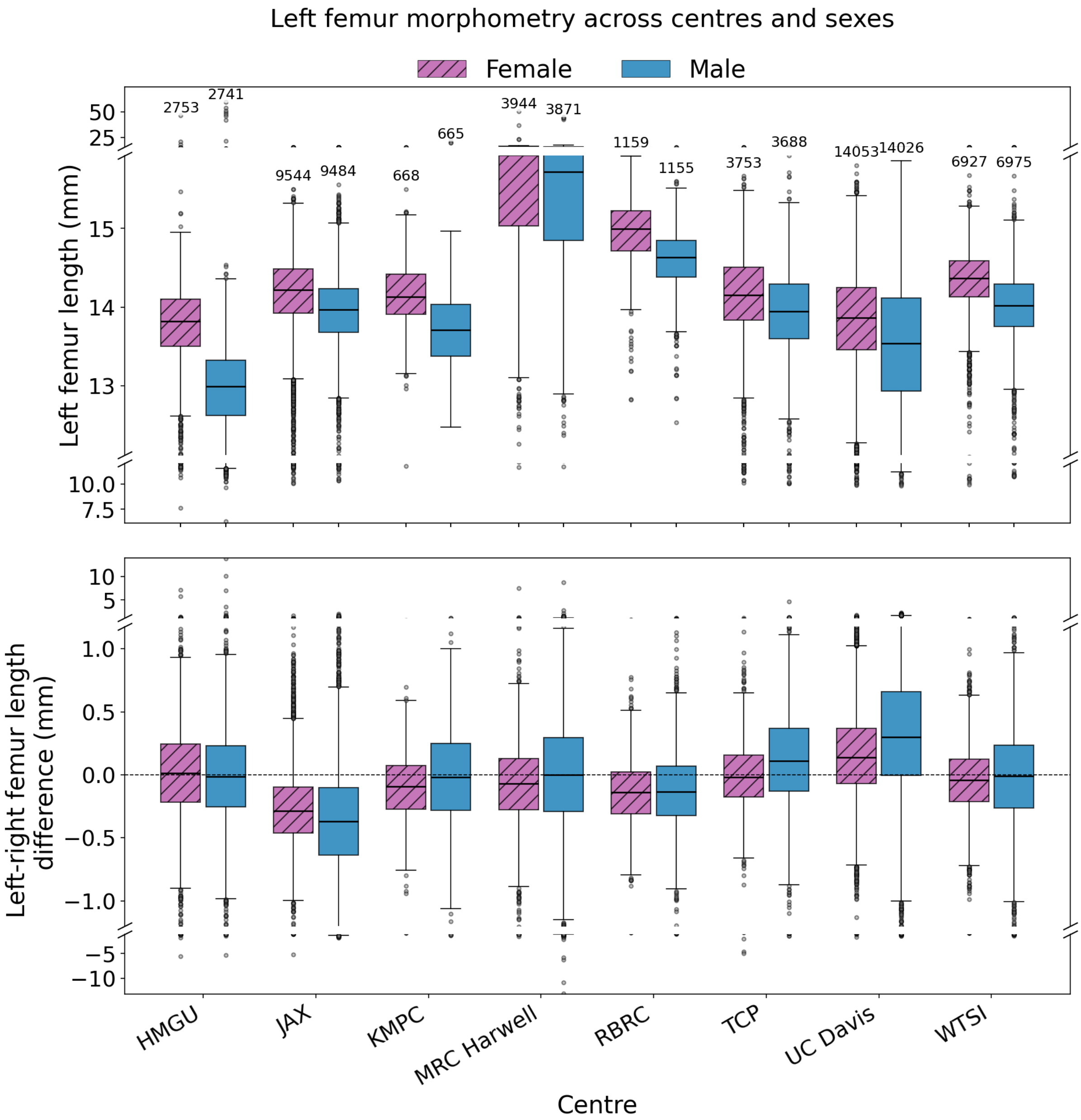


Figure S16: **Supplementary centre- and sex-stratified femur morphometry**. The upper panel shows the distribution of left femur lengths in millimetres for female and male mice within each centre. The lower panel shows the corresponding signed difference between left and right femur length for the same centre/sex strata. Numbers above the upper boxplots indicate the number of radiographs contributing to each centre/sex stratum. Pink boxes denote female mice and blue boxes male mice.

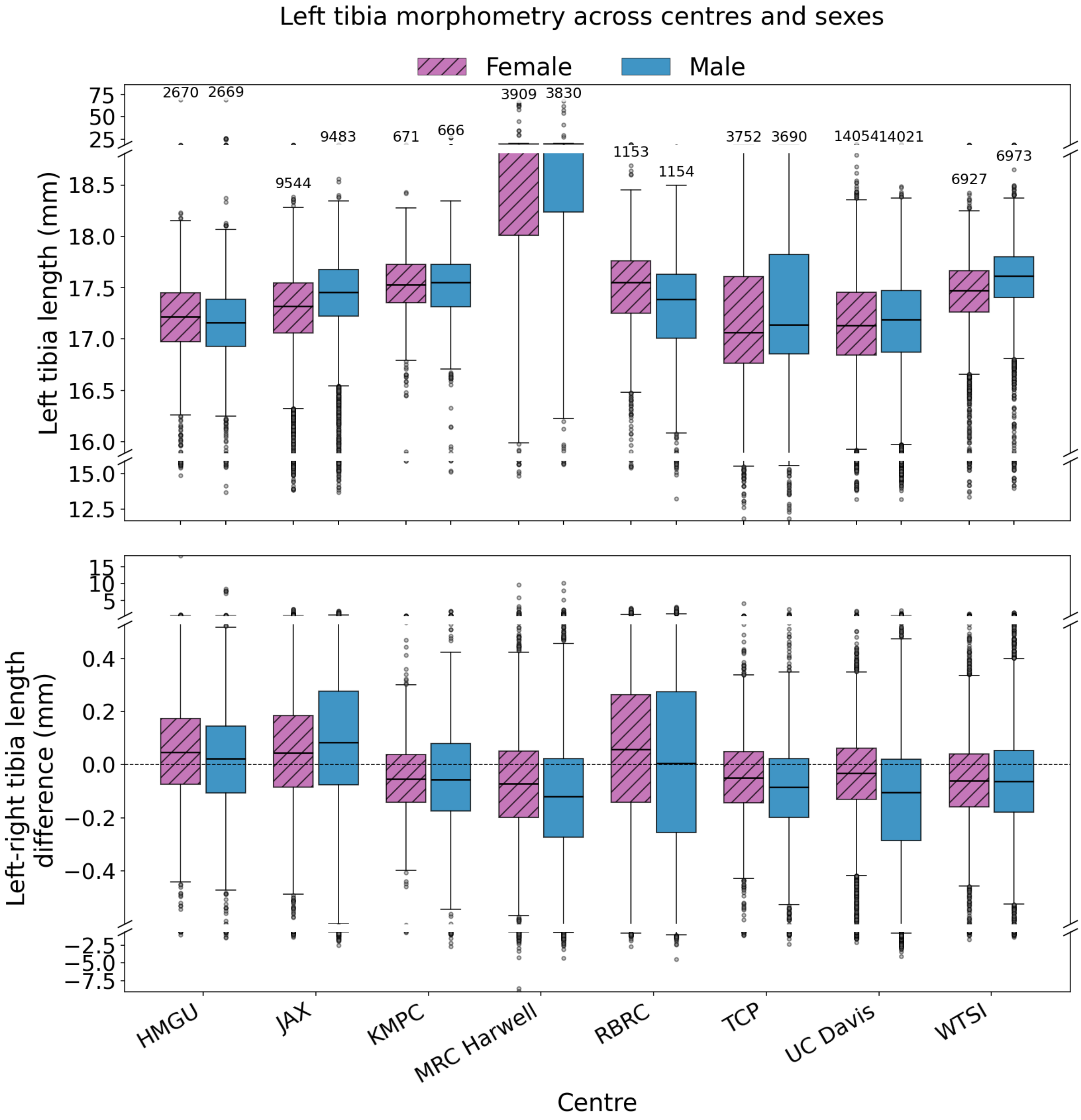


Figure S17: **Supplementary centre- and sex-stratified tibia morphometry**. The upper panel shows the distribution of left tibia lengths in millimetres for female and male mice within each centre. The lower panel shows the corresponding signed difference between left and right tibia length for the same centre/sex strata. Numbers above the upper boxplots indicate the number of radiographs contributing to each centre/sex stratum. Pink boxes denote female mice and blue boxes male mice.

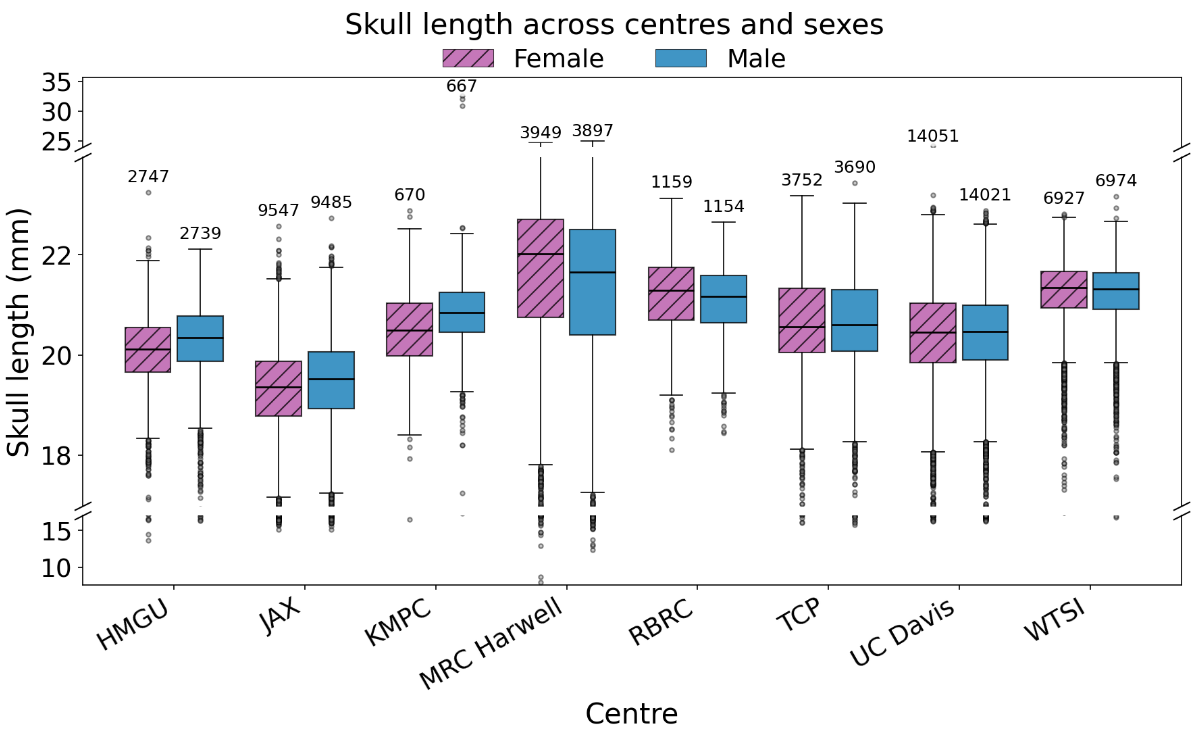


Figure S18: **Supplementary centre- and sex-stratified skull length measurements in millimetres derived from the DeepLabCut-based downstream pipeline**. Numbers above the boxplots indicate the number of radiographs contributing to each centre/sex stratum. Pink boxes denote female mice and blue boxes male mice.

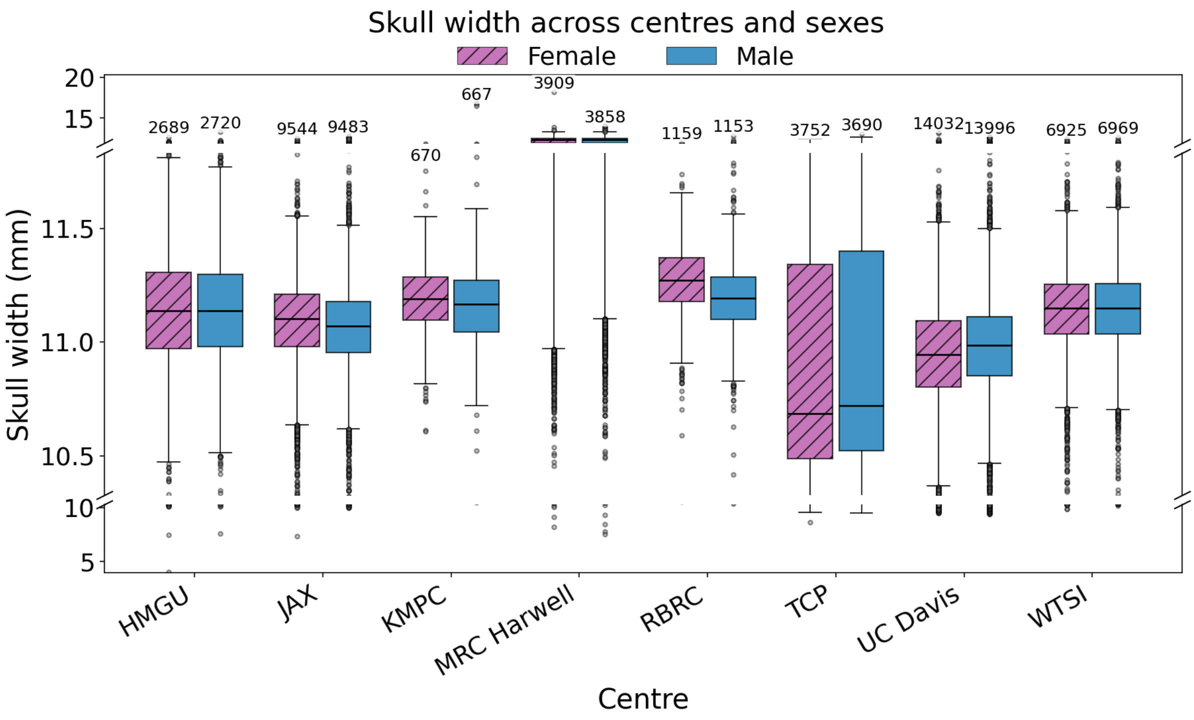


Figure S19: **Supplementary centre- and sex-stratified skull width measurements in millimetres derived from the DeepLabCut-based downstream pipeline**. Numbers above the boxplots indicate the number of radiographs contributing to each centre/sex stratum. Pink boxes denote female mice and blue boxes male mice.

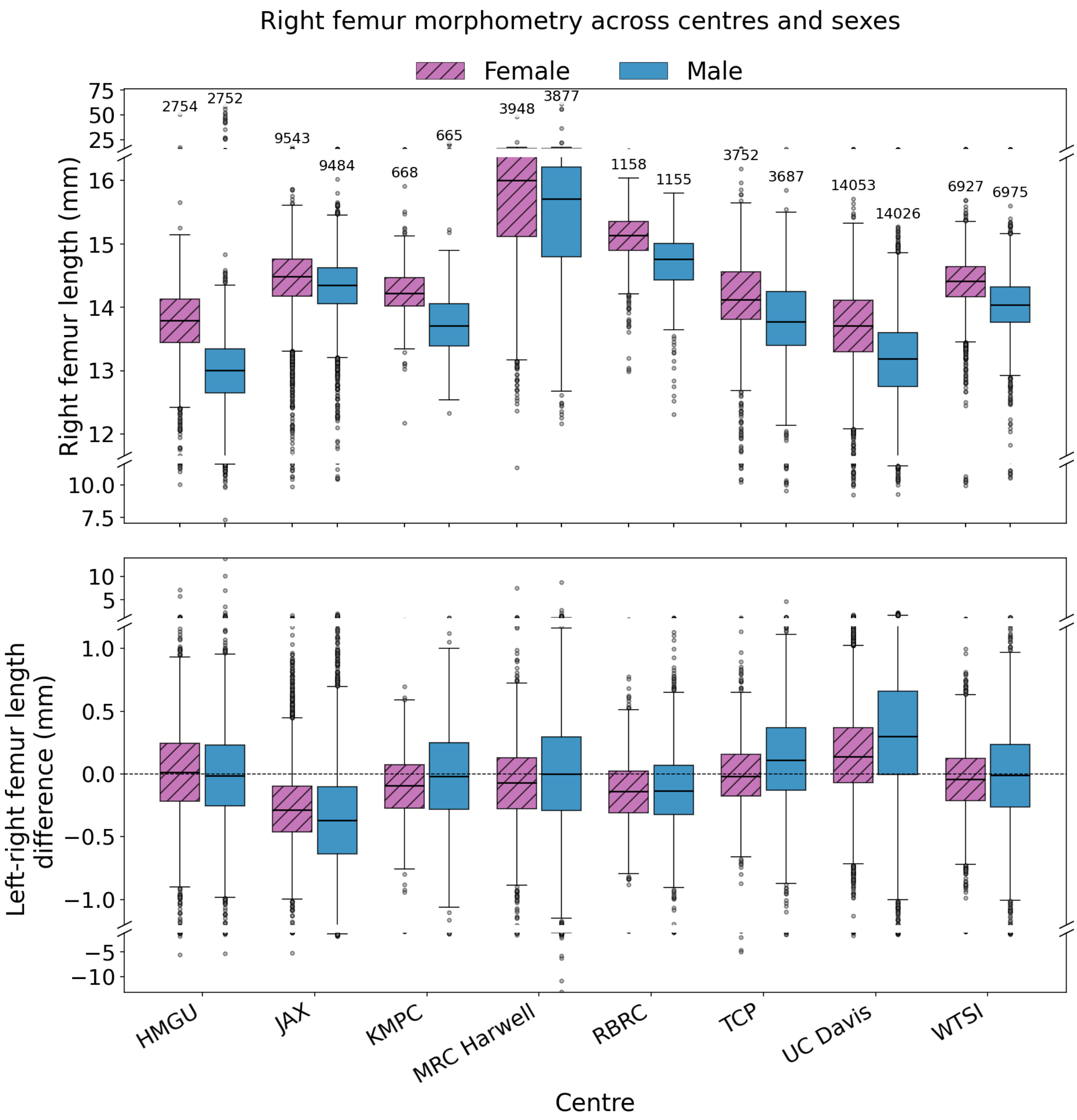


Figure S20: **Supplementary centre- and sex-stratified right femur morphometry**. The upper panel shows the distribution of right femur lengths in millimetres for female and male mice within each centre. The lower panel shows the corresponding signed difference between right and left femur length for the same centre/sex strata. Numbers above the upper boxplots indicate the number of radiographs contributing to each centre/sex stratum. Pink boxes denote female mice and blue boxes male mice.

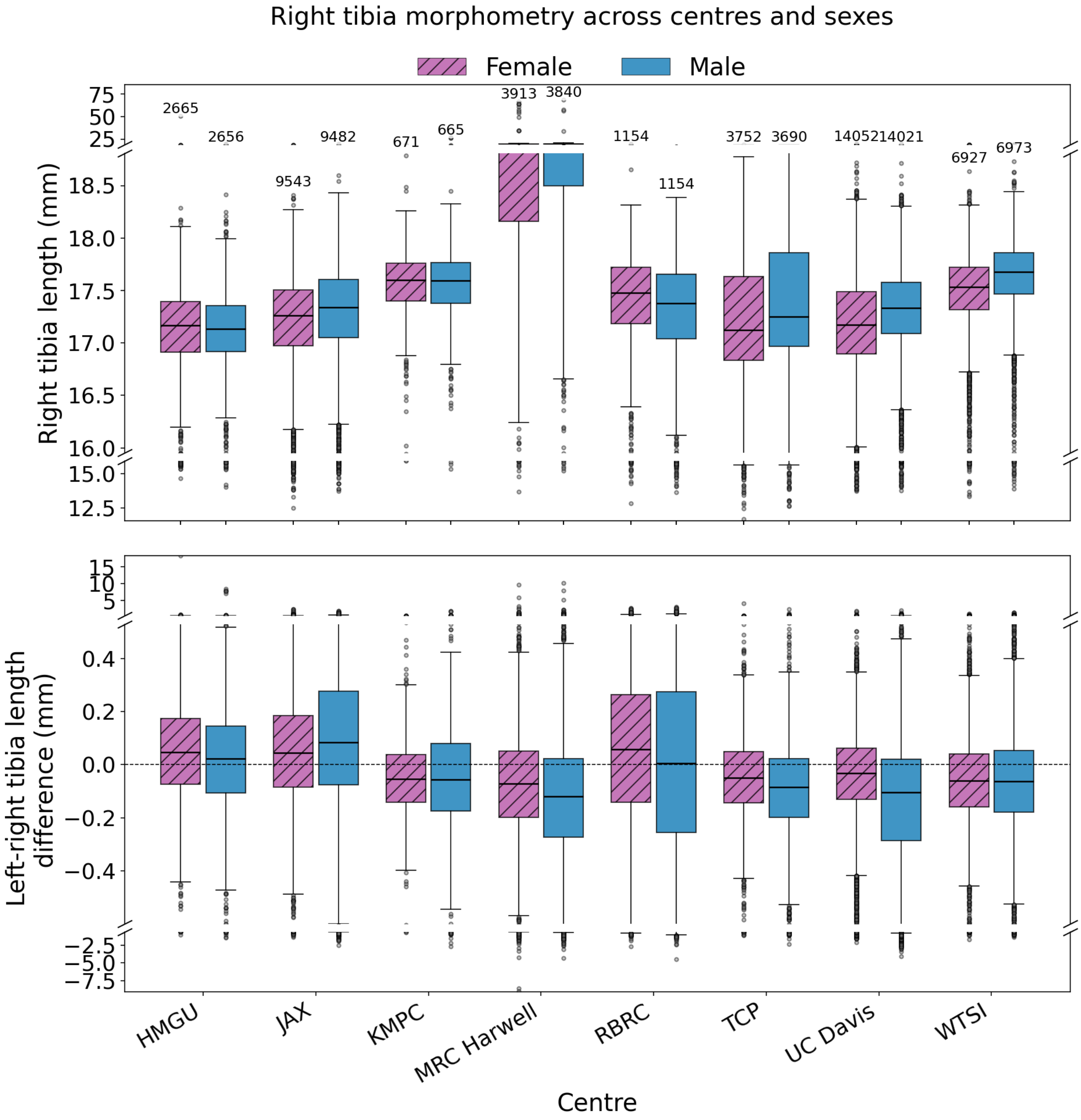


Figure S21: **Supplementary centre- and sex-stratified right tibia morphometry**. The upper panel shows the distribution of right tibia lengths in millimetres for female and male mice within each centre. The lower panel shows the corresponding signed difference between right and left tibia length for the same centre/sex strata. Numbers above the upper boxplots indicate the number of radiographs contributing to each centre/sex stratum. Pink boxes denote female mice and blue boxes male mice.

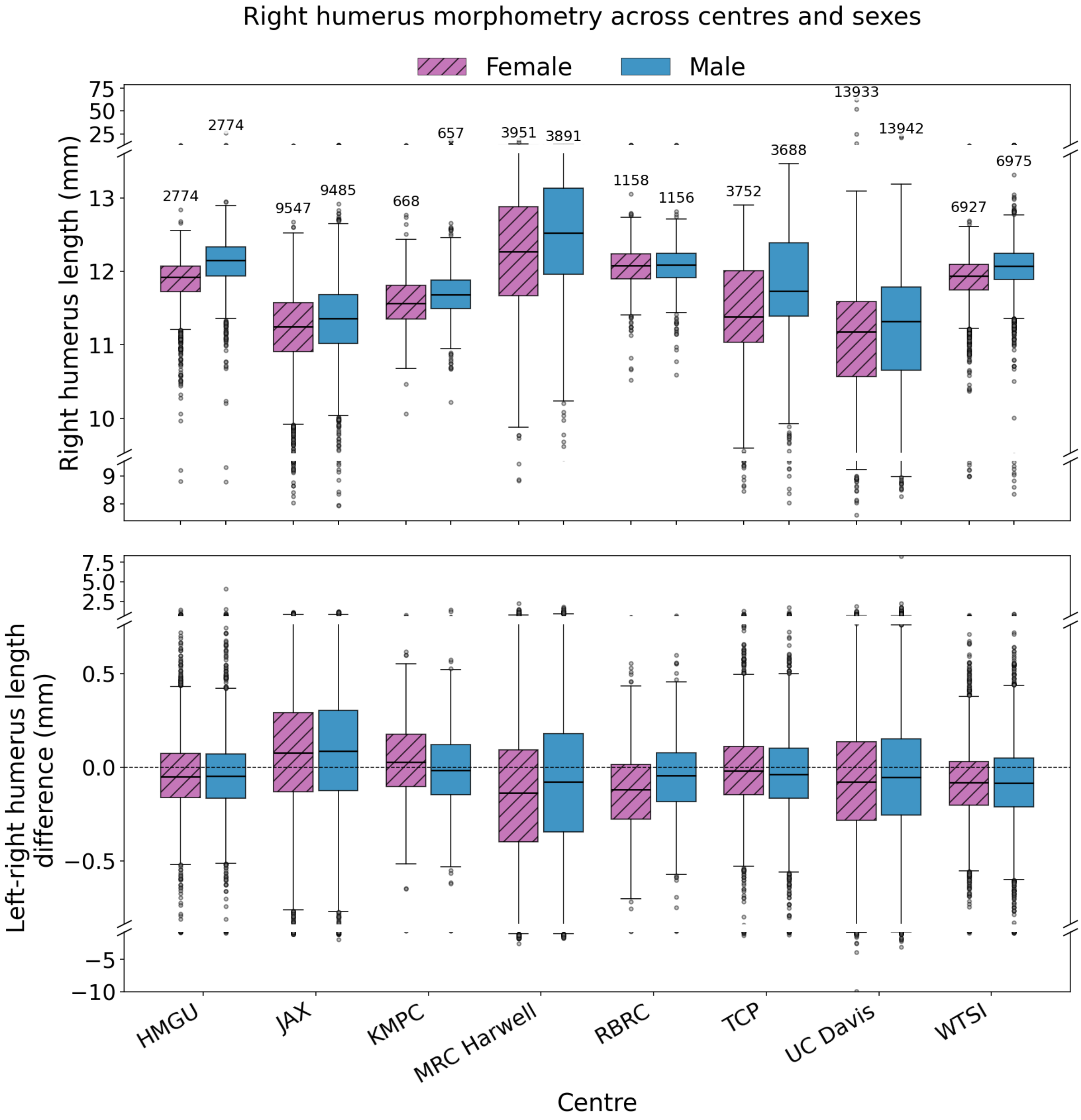


Figure S22: **Supplementary centre- and sex-stratified right humerus morphometry**. The upper panel shows the distribution of right humerus lengths in millimetres for female and male mice within each centre. The lower panel shows the corresponding signed difference between right and left humerus length for the same centre/sex strata. Numbers above the upper boxplots indicate the number of radiographs contributing to each centre/sex stratum. Pink boxes denote female mice and blue boxes male mice.
